# Shared and Unique Neural Codes for Biological Motion Perception in Humans and Macaque Monkeys

**DOI:** 10.1101/2024.08.13.607730

**Authors:** Yuhui Cheng, Yumeng Xin, Xiqian Lu, Tianshu Yang, Xiaohan Ma, Xiangyong Yuan, Ning Liu, Yi Jiang

## Abstract

Throughout evolution, living organisms have honed the ability to swiftly recognize biological motion (BM) across species. However, how the brain processes within- and cross-species BM, and the evolutionary progression of these processes, remain unclear. To investigate these questions, we examined brain activity in the lateral temporal areas of humans and monkeys as they passively observed upright and inverted human and macaque BM stimuli. In humans, the middle temporal area (hMT+) responded to both human and macaque BM stimuli, while the right posterior superior temporal sulcus (pSTS) exhibited selective responses to human BM stimuli. This selectivity was evidenced by an increased feedforward connection from hMT+ to pSTS during the processing of human BM stimuli. In monkeys, the MT region processed BM stimuli from both species, but no subregion in the STS anterior to MT was specific to conspecific BM stimuli. A comparison of these findings suggests that upstream brain regions (i.e., MT) may retain homologous functions across species, while downstream brain regions (i.e., STS) may have undergone differentiation and specialization throughout evolution. These results provide insights into the commonalities and differences in the specialized visual pathway engaged in processing within- and cross-species BMs, as well as their functional divergence during evolution.

## Introduction

Biological motion (BM) refers to the recognizable movement patterns generated by living creatures. These patterns can be detected and recognized even with only a few point lights attached to the head and major joints (Johansson, 1973, 1976). BM signals serve as a remarkable mode of communication, conveying not only the movement status but also the social intention (Johnson, 2006; Manera, Schouten, Becchio, Bara, & Verfaillie, 2010). Therefore, the efficient recognition of BM signals is a cornerstone ability for survival, adaptation and social development. It is under high selective pressure to evolve specialized mechanisms tuned to this ability.

On the one hand, the general ability to identify the movements of organisms in a complex external environment is universally valuable for successful daily-life activities, not limited to any specific species. Numerous studies have demonstrated that both humans and animals can easily identify the point-light BMs from their conspecifics and other species (Blake, 1993; Pinto & Shiffrar, 2009; Troje & Westhoff, 2006; Vallortigara, Regolin, & Marconato, 2005). The intrinsic ability to detect BMs of various species emerges early in life and may be inheritable (Bidet-Ildei, Kitromilides, Orliaguet, Pavlova, & Gentaz, 2014; Simion, Regolin, & Bulf, 2008; Wang et al., 2018). Arguably, these findings collectively indicate that there may exist an evolutionarily ancient and functionally non-specific mechanism, generally tailored to cross-species BM perception (Hirai & Senju, 2020; Troje & Westhoff, 2006).

On the other hand, conspecific BM signals contain a wealth of socially relevant information. Social animals can rely on these signals to interact effectively with each other and engage in collaborative group activities in order to satisfy their intrinsic social desires and maintain social bonds (Thornton, 2012). Therefore, from the perspective of social importance, perceptions of conspecific and non-conspecific BM are not homogenous. Plenty of studies have reported that humans are highly sensitive to conspecific information, as evidenced by quicker and more accurate discrimination of human stimuli compared to non-human animal stimuli (Boucart et al., 2016; New, Cosmides, & Tooby, 2007; Pinto & Shiffrar, 2009; Stein, Sterzer, & Peelen, 2012). These studies imply that living creatures may have evolved a specific mechanism to recognize conspecific BM. This mechanism is greatly favored by natural selection for effective social engagement.

Accordingly, the neural mechanisms underlying BM perception may be theoretically characterized as species-general and species-specific. However, it is not well understood whether there are corresponding neural substrates that respectively underscore the species-general and species-specific BM processing, or how these neural mechanisms develop and evolve across subjects with the same ancestor. Addressing these issues requires a cross-species design, in which humans and macaque monkeys are engaged in a parallel study employing an identical factorial design to investigate their brain activities associated with BM perception. Such design allows for the comprehensive exploration and direct comparison of the neural mechanisms involved in processing within- and cross-species BMs between humans and monkeys, who share a common ancestor and have the closest evolutionary relationship.

For humans, research has demonstrated that processing BM information activates multiple regions in the superior temporal sulcus (STS), middle temporal cortex (MT), fusiform gyrus (FFG), portions of the frontal and parietal cortex (Grosbras, Beaton, & Eickhoff, 2012; Grossman & Blake, 2002; Peelen, Wiggett, & Downing, 2006), as well as the left cerebellum and some subcortical brain regions (H. F. Chang, Ban, Ikegaya, Fujita, & Troje, 2018; Pavlova et al., 2017; Sokolov et al., 2012; Sokolov et al., 2018). Among these regions, the most consistently involved are the human middle temporal area (hMT+) and the posterior superior temporal sulcus (pSTS) (Grosbras et al., 2012). Similar brain regions selectively sensitive to BM stimuli are found in the temporal cortex of monkeys, which includes the monkey MT complex, anterior regions of the fundus of the STS, such as the lower superior temporal region (LST), and the dorsal bank of the STS, such as the TEO region (Huk, Dougherty, & Heeger, 2002; Jastorff, Popivanov, Vogels, Vanduffel, & Orban, 2012; Nelissen, Vanduffel, & Orban, 2006; Russ & Leopold, 2015). Recently, it has been proposed that social dynamic information (including BMs and moving faces) in humans and monkeys mainly transmits in a third visual pathway from the primary visual cortex via MT into STS (Pitcher & Ungerleider, 2021). The brain regions in the upstream and downstream of this pathway (e.g., MT and STS) are selectively tuned to hierarchical aspects of the visual signals, from simple motion patterns (Born & Bradley, 2005) to action intentions (Isik, Koldewyn, Beeler, & Kanwisher, 2017; Jastorff et al., 2012).

Based on previous findings, the current study focuses on these two key brain regions (MT and STS) in the third visual pathway. The aim of this study is twofold. The first is to delineate whether within- and cross-species BMs, which primarily differ from each other in social aspects, may activate the two brain areas differently. The second is to further examine whether humans and non-human primates (i.e., monkeys), who share a common ancestor and have the closest evolutionary relationship, process the within- and cross-species BMs in a similar or different manner.

To this end, the current study used functional magnetic resonance imaging (fMRI) to measure brain activity in both human and monkey participants as they viewed human and macaque BM stimuli. Following other studies, we compared the brain responses to upright and inverted versions of the BM stimuli. This comparison allowed us to examine the neural correlates of animal movements while minimizing the influence of low-level visual factors (H. F. Chang et al., 2018; Troje & Westhoff, 2006; Wang et al., 2022). We collected brain signals from the whole brain and from the predefined regions of interest (ROIs) in both humans and monkeys. We then applied univariate analysis, multivoxel pattern analysis (MVPA) within and across species, as well as brain connectivity analysis to identify the respective brain regions underlying the species-general and species-specific BM perception.

## Results

### Human fMRI Experiments

#### Behavioral Performance

During the fMRI scan, human participants viewed the upright and inverted versions of point-light BM stimuli. These stimuli were derived from different species: human BM or macaque BM (see Figure 1). The participants were asked to perform a task involving the detection of color changes in the fixation point. This task was designed to ensure that they maintained their attention on the display throughout the scan. The mean accuracy of the participants across all types of BM stimuli was 84.46 ± 22.55%. Importantly, there were no significant differences in accuracy between the upright and inverted versions of each type of BM stimuli (*ps* > 0.299 for all comparisons), indicating that the participants were able to uniformly focus their attention on both the upright and inverted versions across different types of BM stimuli.

**Figure 1.**
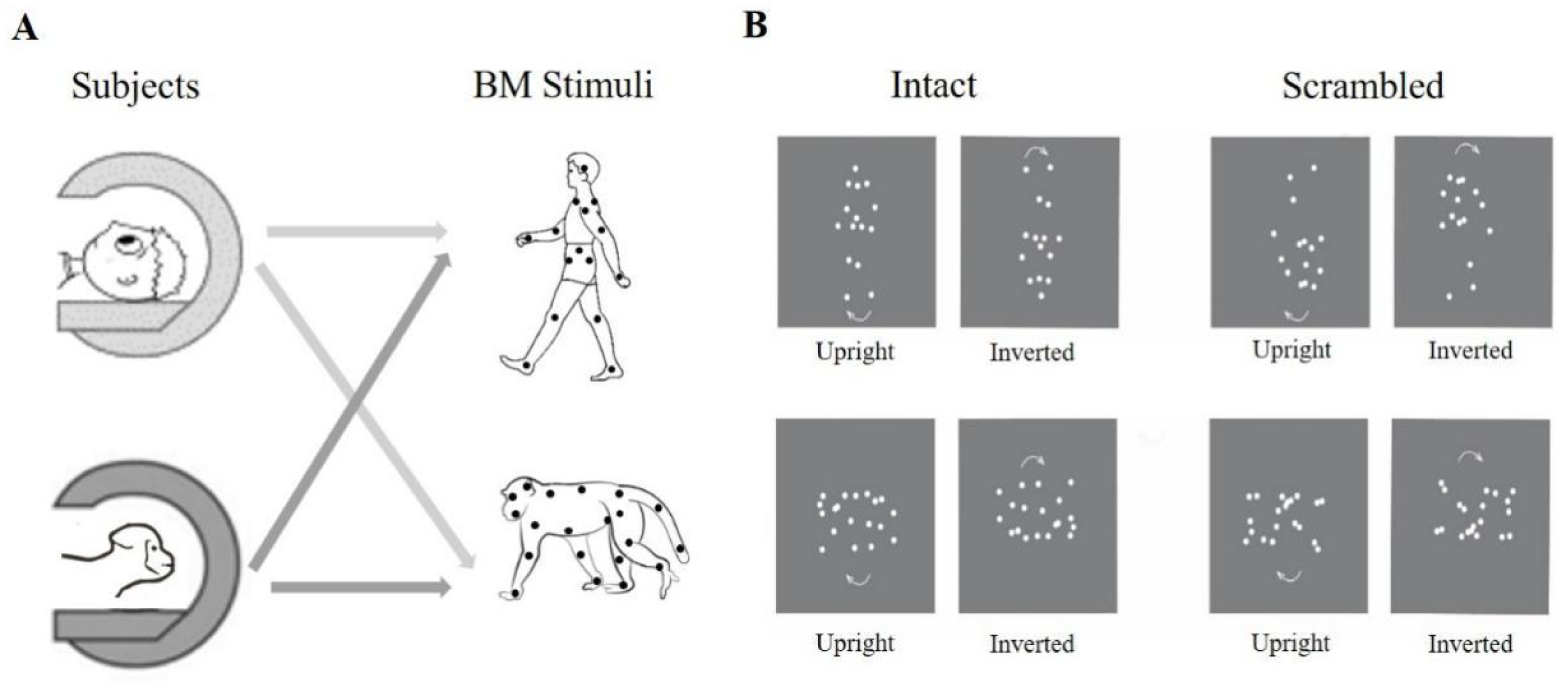
Schematic illustration of experimental design and sample stimuli. (A) Human and monkey participants viewed both human and macaque BM stimuli during the fMRI scans. (B) Intact human BM, scrambled human BM, intact macaque BM, and scrambled macaque BM stimuli were used in the human and monkey fMRI experiments, including their upright and inverted versions. Arrows indicated the motion direction and were not presented in the actual experiments.

#### Univariate Results

Firstly, the whole-brain analysis confirmed the engagement of the MT and STS in BM perception. The results are illustrated in Figure 2A (refer to Table S1 for more details). The contrast between upright and inverted BM showed that the intact human BM stimuli elicited increased neural activities in the right middle temporal gyrus extending to the right posterior superior temporal sulcus (pSTS), while the intact macaque BM stimuli only elicited stronger brain activation in the middle occipital gyrus (extending to the middle temporal gyrus). A conjunction analysis was further performed to provide clues about the human brain activation in response to both within- and cross-species BM stimuli. The results showed that the brain activation for intact human and macaque BMs overlapped in the right MT (Figure 2B, peak at x = 42, y = -76, z = 6, 55 voxels, with a voxelwise FDR-corrected threshold of *p* < 0.05). The whole-brain findings demonstrated that the MT area is capable of representing cross-species BM information, whereas the right pSTS is specifically activated by the intact human BM (i.e., the same species). The same contrast did not reveal any significantly increased brain activation for the scrambled human BM stimuli except for a stronger activation in the lateral occipital cortex for the scrambled macaque B stimuli. Given that the scrambled BM stimuli failed to elicit comparable activation to the intact BM stimuli in the current univariate and subsequent multivariate analyses (available in supplementary material B), we did not further discuss them in the main text and focused solely on the results obtained from the intact BM stimuli.

**Figure 2.**
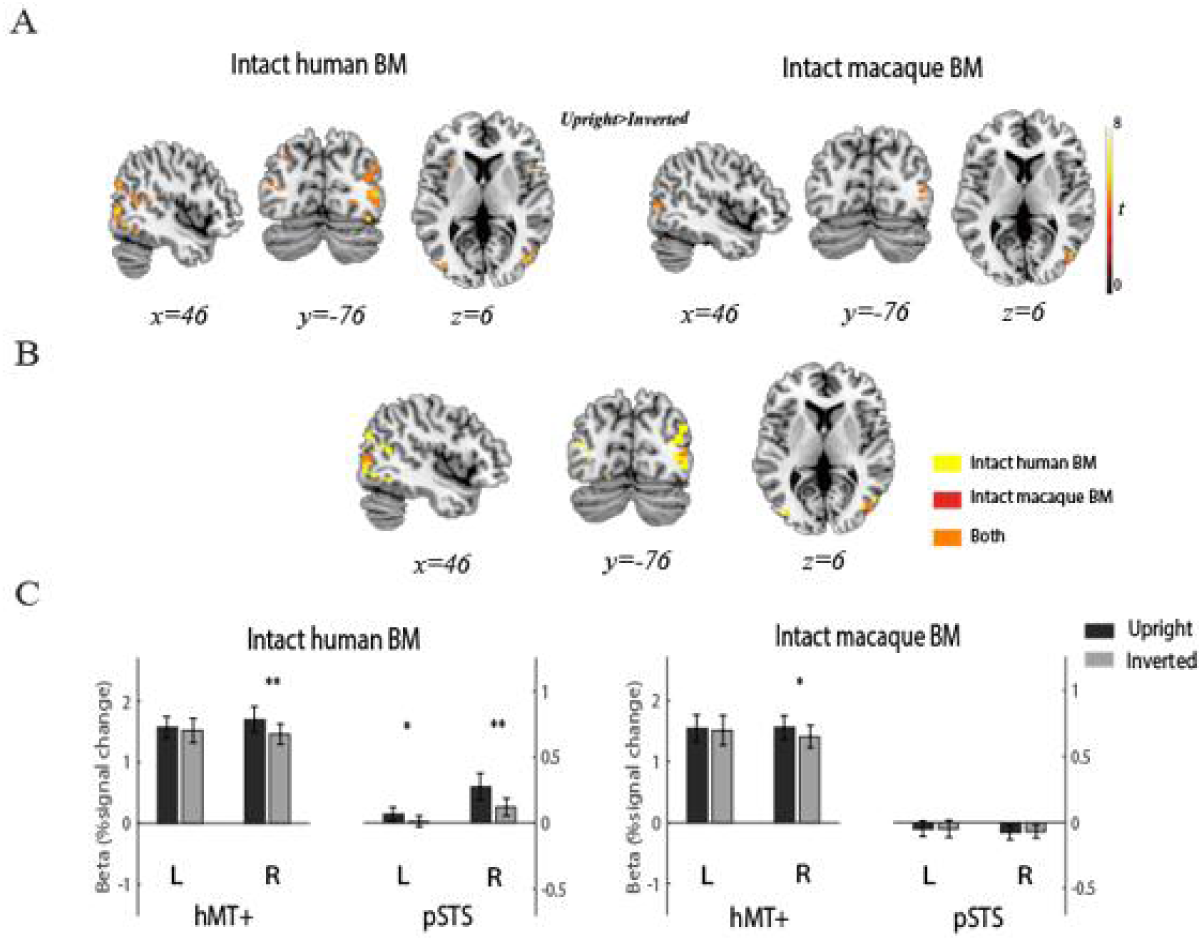
Group-level fMRI activation maps of human whole-brain and ROI results. (A) Regions depicting areas of activation to the upright versus inverted intact human and macaque BM stimuli with an FDR-corrected threshold of *p* < 0.05 at the cluster level and *p* < 0.001 uncorrected at the voxel level (the minimum cluster size > 10 voxels). The color bar indicates *t* values. (B) The common region identified by the conjunction analysis of intact human BM and intact macaque BM. (C) Mean beta values across participants, in each individually defined left and right hMT+ and pSTS, in response to upright versus inverted BM, respectively. Error bars denote standard errors of the mean. \**p* < 0.05, \*\**p* < 0.01.

To further examine the species-general and species-specific BM processing in MT and STS, we defined these two brain regions using independent localizer tasks (i.e., hMT+ and pSTS; see the Method section, Table S2 and Figure S1 for more details), and performed 2 (Hemisphere: left vs. right) × 2 (Orientation: upright vs. inverted) repeated measures analysis of variance (ANOVA) on them for each type of BM stimuli. The results are shown in Figure 2C (please refer to Table S3 for detailed statistics). In brief, we found that the intact human BM stimuli produced significant main effects of Orientation and significant interactions in both ROIs (hMT+: *F*_1,17_ > 6.16, *p* < 0.024; pSTS: *F*_1,17_ > 6.36, *p* < 0.022). The post-hoc analysis revealed the upright intact human BM, relative to their inverted counterparts, significantly activated the right hMT+ (right: *t*_17_ = 3.56, *p* = 0.002) and bilateral pSTS (left: *t*_17_ = 2.63, *p* = 0.017; right: *t*_17_ = 3.89, *p* = 0.001). For the intact macaque BM stimuli, only a significant interaction effect was observed in the hMT+ (*F*_1,17_ = 11.61, *p* = 0.003). Similarly, the upright intact macaque BM, compared with the inverted one, selectively activated the right hMT+ (*t*_17_ = 2.79, *p* = 0.013). Conversely, no significant effect was observed in the pSTS (*F*s < 2.29, *ps* > 0.149). In sum, the ROI results again confirmed the whole-brain results, showing that the hMT+ is generally dedicated to cross-species BM perception, whereas the pSTS, especially in the right hemisphere, is specifically dedicated to same-species BM perception.

#### Multivariate Results

Using MVPA, we further tested whether the brain activation pattern in the hMT+ and pSTS differentially represents within- and cross-species BM stimuli. First, we trained the classifier to categorize the upright and inverted human/macaque BM stimuli based on the activation pattern of all voxels in the predefined hMT+ and pSTS, respectively. We then tested the classification accuracy on the leave-out sample. We found that the classification accuracies in both ROIs were significantly above chance level for the intact BM displays (see Figure 3A and Table S4). This result suggests that, in addition to the hMT+, the activity pattern in the right pSTS region can recognize same-species BM stimuli (human), whereas the left pSTS region can recognize BM stimuli from other species (macaque). However, it remains unclear whether these regions can accurately decode the common BM information shared across species.

**Figure 3.**
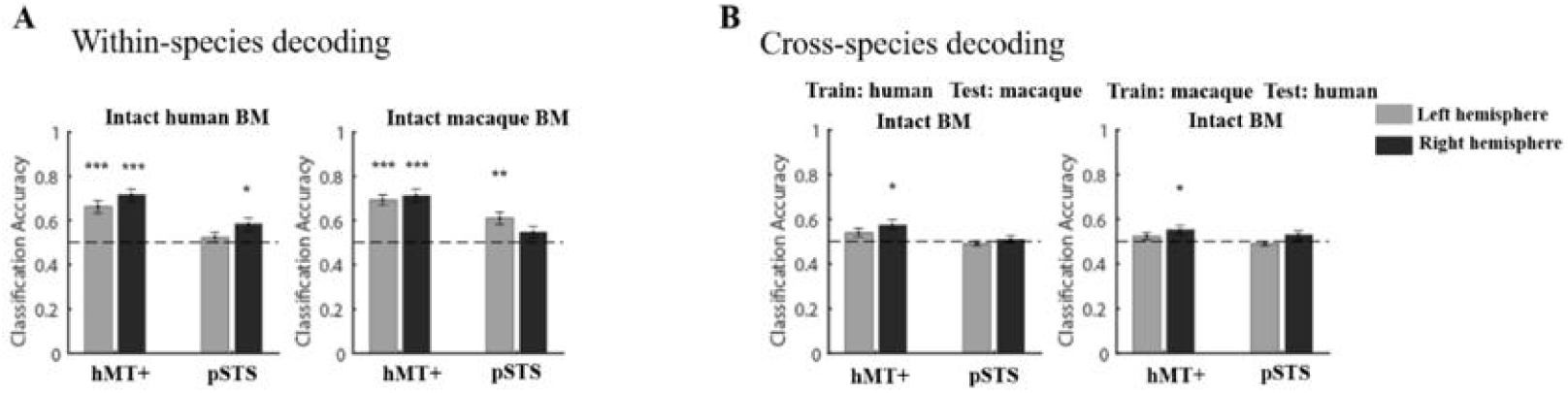
The human MVPA results. The classification accuracies of within-species (A) and cross-species (B) decoding of upright and inverted BM stimuli in the hMT+ and pSTS. The horizontal dashed line represents the chance level (50%). Error bars denote standard errors of the mean. \**p* < 0.05, \*\**p* < 0.01, \*\*\**p* < 0.001.

To answer this question, we trained the classifier to categorize the intact upright and inverted BM stimuli from one species (e.g., human BM) and tested its classification accuracy on the BM stimuli from the other species (e.g., macaque BM). The results revealed that, irrespective of the species of the stimuli, the intact upright and inverted BM stimuli could be well discriminated in the right hMT+ (train human BM and test macaque BM: *t*_17_ =3.24, *p* = 0.019; train macaque BM and test human BM: *t*_17_ = 2.86, *p* = 0.044), but not in the pSTS (*ts* < 1.45, *ps* >0.631), as illustrated in Figure 3B and Table S5. The cross-species MVPA clearly demonstrated that hMT+ can generally decode cross-species BMs, whereas the pSTS, especially in the right hemisphere, specifically decodes within-species BMs.

#### Psychophysiological Interaction and Dynamic Causal Modeling Results

Having revealed the species-general and specific properties of BM processing in the hMT+ and pSTS, it is intriguing to explore how the within- and cross-species BM information is transmitted from MT to STS in the third visual pathway using functional connectivity analysis. We first implemented an exploratory psychophysiological interaction (PPI) analysis using the bilateral hMT+ as a search seed. The results showed that the right pSTS (x = 64, y = -36, z = 6; *p* < 0.05, corrected for multiple comparisons) was co-activated with the bilateral hMT+ when the intact human BM stimuli were displayed. Other regions functionally connected with the bilateral hMT+ included the left superior frontal gyrus, the bilateral cingulate gyrus, the left anterior cingulate, the left supramarginal gyrus, and the right middle temporal gyrus (see Figure S2 and Table S6). However, no significant clusters with the predefined threshold (*p* value) were revealed when the same PPI analysis was performed for the intact macaque BM stimuli.

To further evaluate the directed functional connectivity between the hMT+ and pSTS, we conducted a dynamic causal modeling (DCM) analysis. First, we assumed that the driving visual inputs, including the upright and inverted BM stimuli, enter the system through the hMT+ (C-matrix), and that the intrinsic connections between the hMT+ and pSTS are all bidirectionally fixed (hMT+ ↔ pSTS; A-matrix). We then systematically assessed the connections mediated by BM processing (upright > inverted) across the following four models: (1) no modulation, (2) forward modulation, (3) backward modulation, and (4) bidirectional modulation (Figure 4A). The results indicated that the “bidirectional modulation” had the highest posterior probability for the intact human BM stimuli, while the “forward modulation” had the highest posterior probability for the intact macaque BM stimuli (Figure 4B). Paired sample t-tests showed that, for the best models, the driving inputs of the upright BM stimuli were significantly or marginally significantly stronger than those of the inverted ones for both the intact human (*t*_17_ = 2.12, *p* = 0.047) and macaque BM perception (*t*_17_ = 1.82, *p* = 0.087). However, for the intact human stimuli only, the modulatory effect of the upright BM was significantly stronger than that of the inverted BM on the forward connection from hMT+ to pSTS (*t*_17_ = 4.76, *p* < 0.001), but not on the backward connection: *t*_17_ = -0.02, *p* = 0.981). For the intact macaque stimuli, the modulatory effect on the forward connection was not significant (*t*_17_ = 1.25, *p* = 0.229) and no backward connection was found. Taken together, these PPI and DCM results consistently demonstrated that the functional connection between hMT+ and pSTS, especially the feedforward one, is indispensable for within-species BM perception. Therefore, it is probable that the cross-species BM information decoded in the hMT+ is not transmitted to the pSTS as fluently as the within-species BM information.

**Figure 4.**
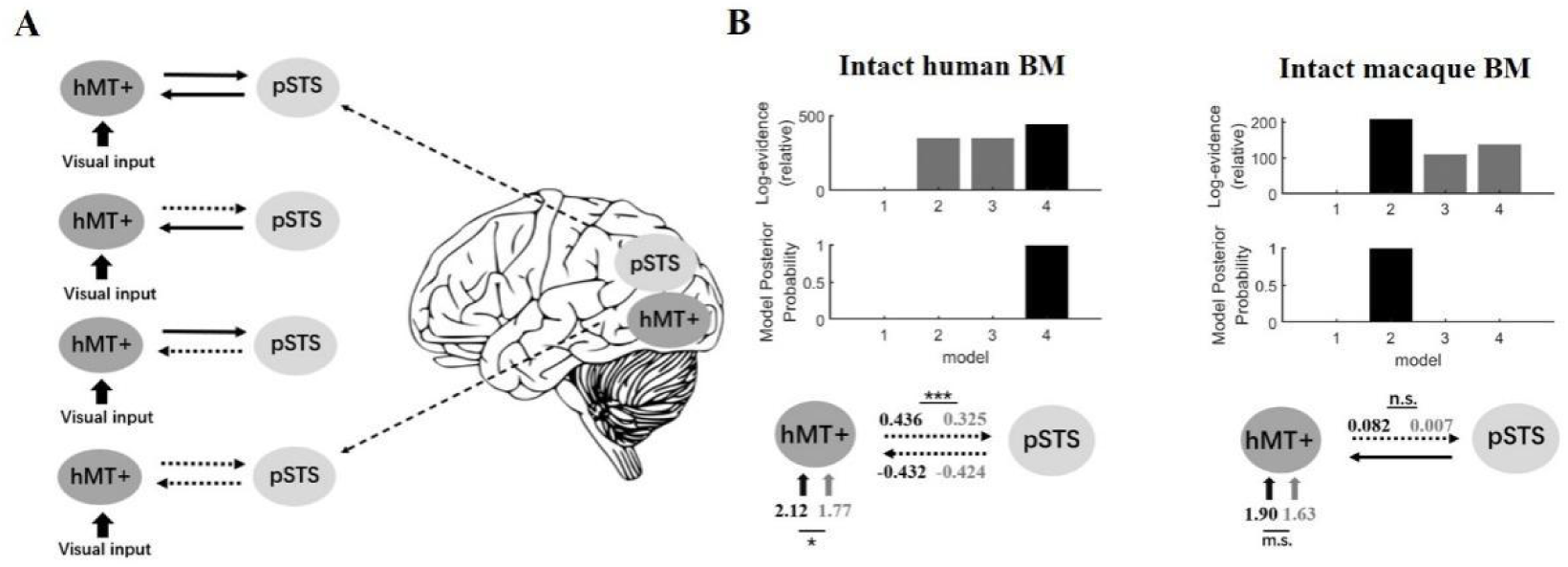
Models and results of DCM. (A) The four DCM models between hMT+ and pSTS are presented from top to bottom, in the following order: ‘no modulation’, ‘forward modulation’, ‘backward modulation’, and ‘bidirectional modulation’. The solid bold arrows indicate the driving visual inputs. The horizontal black arrows indicate the intrinsic connections between hMT+ and pSTS. The horizontal dotted arrows indicate the intrinsic connections assumed to be modulated by BM perception. (B) The DCM results for intact human and macaque BM perception (upright versus inverted). The top panels show the log-evidence and the posterior probability for all models. The bottom panels show the parameters for driving inputs and modulatory effects (expressed in Hz). The black and the gray arrows separately indicate the parameters from the upright and inverted BM. *n.s.,* not significant, *m.s.,* marginally significant, ∗*p* < 0.05, ∗∗∗*p* < 0.001.

Finally, since the pSTS ROIs were identified separately for the human and macaque BM stimuli in the localizer task and did not fully overlap with each other (see the Method section), one might argue that the species-specific properties of the pSTS could be attributed to this ROI definition. To rule out this possibility, we redefined a combined pSTS using the human and macaque BM stimuli together as a whole and re-conducted the ROI analysis, MVPA analysis, and functional connectivity analysis on this combined ROI. The results replicated the main conclusion above, and more details can be found in Supplemental Material section B. Therefore, although the right hMT+ enables cross-species and shared BM representation, the downstream region pSTS selectively represents the within-species BMs.

### Monkey fMRI Experiments

#### Behavioral Performance

Similar to the procedure for human participants, monkeys passively observed different types of BM stimuli. They were required to maintain fixation on a central square on the screen throughout the entire experimental session and received a liquid reward. Their eye position was tracked to ensure that there was no significant difference between the fixation percentages of the upright and inverted conditions for each type of BM stimuli (*ps* > 0.058). This indicates that monkeys uniformly focused their attention on both the upright and inverted versions across different types of BM stimuli.

#### Univariate Results

Similarly, using univariate whole-brain analysis with the contrast of upright > inverted BM, we primarily searched for brain activation in the third visual pathway (e.g., MT and STS) that underlies within- and cross-species BM perception in the monkey subjects (Figure 5A). The upright intact macaque BM stimuli elicited significantly stronger neural activities in the MT, V4t, and V4 areas across all three monkeys than the inverted counterparts did, while the intact human BM stimuli produced a similar but less pronounced activation in these regions. These results demonstrated that the monkey MT and neighboring brain regions were able to encode both same-species and cross-species intact BM. Indeed, the conjunction analyses confirmed this result (Figure 5A). However, we did not find any other significant activations in the STS regions anterior to MT with the same contrast across the three monkeys.

**Figure 5.**
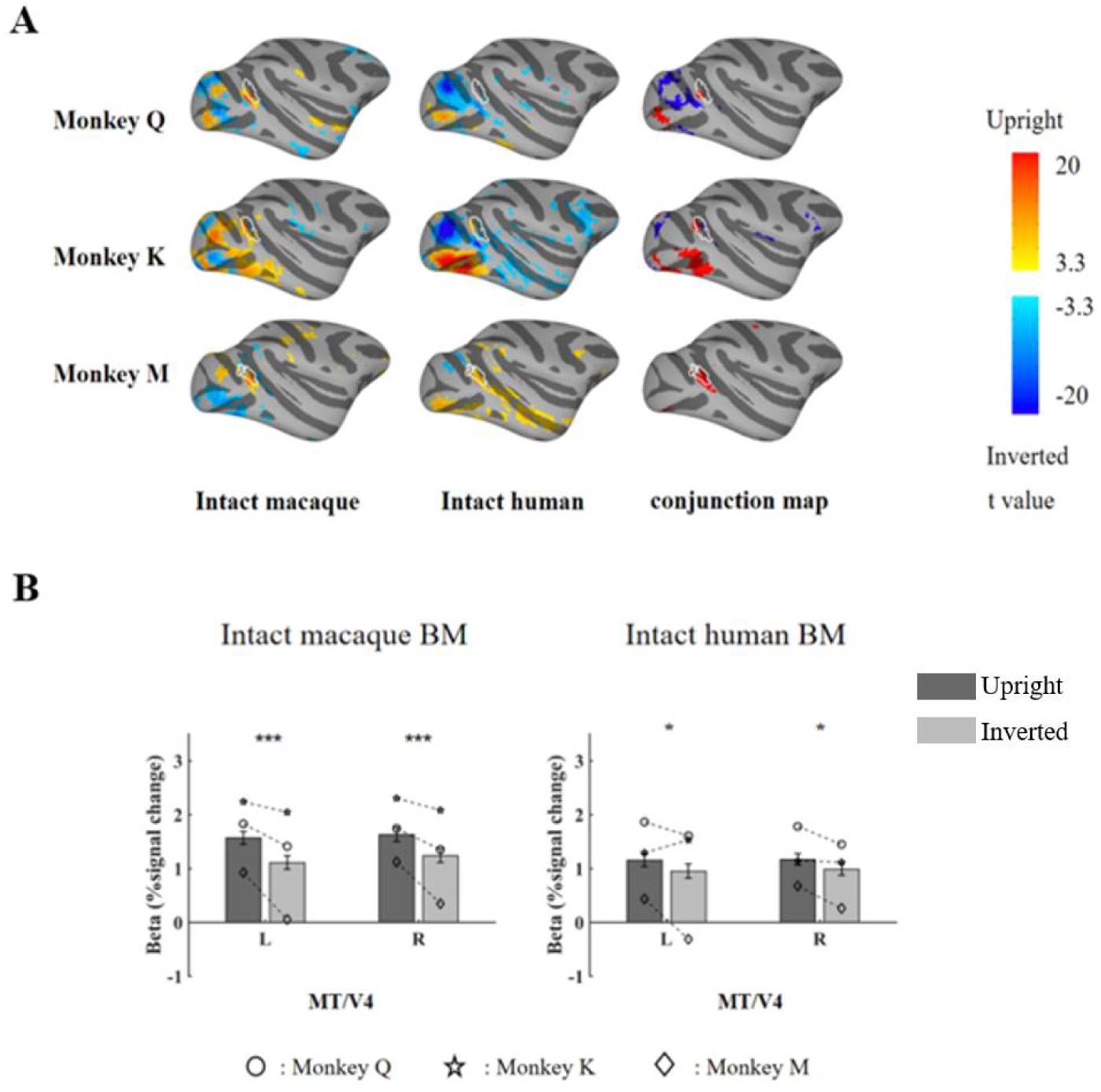
Monkey whole-brain analyses and ROI results. (A) Brain activation maps of three monkeys for the intact BM stimuli (upright versus inverted), shown on lateral views of the inflated cortex of the right hemisphere. Panels from top to down represent the results of monkeys Q, K, and M, respectively. Panels from left to right show results from the intact macaque and human BM stimuli (uncorrected *p* < 0.001, FDR corrected *p* < 0.05), and the conjunction maps based on them (FDR corrected *p* < 0.05), respectively. The borders of the MT/V4 ROI are encircled by white lines. (B) Results of the ROI analysis in the MT/V4. Error bars indicate standard errors of the mean. The symbols (circle, star, and diamond) within individual bars represent results from monkeys Q, K, and M, respectively. \**p* < 0.05; ***, *p* < 0.001.

Following the analysis protocol of the human experiment, we further examined the above results using ROI analysis (see the Method section along with Figure S4, Table S7, and Table S8 for details). We replicated the whole-brain results in the MT/V4 area by performing a 2 Hemisphere × 2 Orientation Linear Mixed Models (LMMs) with run, session, and monkey as random factors. In short, for the intact macaque BM stimuli, significant main effects of Orientation were found (*F*_1,153_*=* 59.08, *ps* < 0.001), with stronger activation elicited by the upright BM stimuli compared to the inverted ones in the bilateral MT/V4 (left: *t*_153_ = 5.61, *ps* < 0.001; right: *t*_153_ = 5.26, *ps* < 0.001). A similar activation pattern was also found for the intact human BM stimuli (*F*_1,168_ = 11.16, *p* = 0.001), with significant activation elicited by the upright BM stimuli compared to the inverted ones in the bilateral MT/V4 (left: *t*_168_ = 2.27 *p* = 0.025; right: *t*_168_ = 2.46, *p* = 0.015). Note that, in both the human and monkey BM localizers, we found no clusters in the STS responded more strongly to the upright intact BM stimuli than the inverted scramble ones (except those areas close to and overlapped with the MT/V4 ROI in the monkey BM localizer, see Figure S4). Detailed information can be found in Table S9. In sum, echoing the whole-brain results, the monkey ROI results indicated the general abilities to process BM from different species in the MT region, while the STS activation pattern in monkeys differs from that of humans.

#### Multivariate Results

Using within-species MVPA, we measured whether the upright and inverted BM stimuli from the same species could be discriminated in the MT/V4 (Figure 6A). The classification accuracies in the bilateral MT/V4 were significantly higher than chance level for both the intact macaque BM and human BM (see Table S10 for the precise statistical values). Furthermore, we performed MVPA across the BM stimuli from different species (Figure 6B and Table S11). The classification accuracies in the bilateral MT/V4 were significantly higher than chance level, either by training on the human stimuli and testing on the macaque stimuli (left: 74.04 ± 4.30%, right: 81.73 ± 3.79%, *ps* < 0.001), or by training on the macaque stimuli and testing on the human stimuli (left: 64.04 ± 4.49%, right: 71.93 ± 4.21%, *ps* < 0.004). Overall, the monkey MVPA results indicated that the MT/V4 can encode the intact BM stimuli both within-species and cross-species, suggesting a species-general representation of BM in this brain region, similar to the human results of the hMT+.

**Figure 6.**
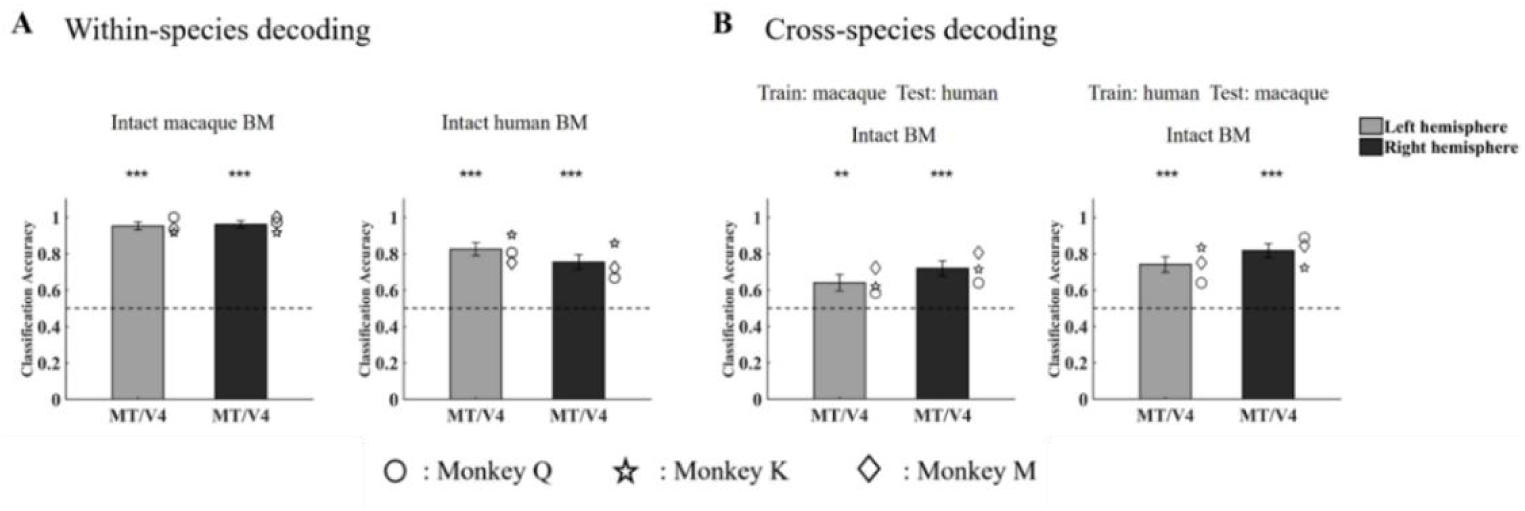
The monkey MVPA results. The classification accuracies of within-species (A) and cross-species (B) decoding of upright and inverted BM stimuli in the MT/V4. Error bars indicate standard errors. The symbols (circle, star, and diamond) on the right side of the individual bar represent results from monkeys Q, K, and M, respectively. The black dashed lines indicate chance level (50%). †, marginally significant, \*\**p* < 0.01, \*\*\**p* < 0.001.

#### Supporting analysis and results

Note that in monkeys, we did not find any clusters at a location similar to the human pSTS using univariate analyses. To further investigate the engagement of monkey STS in BM perception, we conducted the following supporting analysis (see Supplemental Material section D for detailed protocols). Firstly, we attempted to identify clusters in the monkey STS based on the multivariate results. Briefly, we performed whole-brain searchlight analyses for within-species and cross-species classifications (Figures S9 and S10). Based on the results in humans, we conjugated areas that could make successful within-species classifications and those that could not perform successful cross-species decoding (Figure S11). We did find one consistent region anterior to MT/V4 in the posterior inferotemporal cortex (TEO) across the three animals, which exhibited similar performance to the pSTS in humans in the multivariate analyses (Figure S13). Interestingly, we did not observe any significant activation differences between the upright and inverted versions across all four types of BM stimuli (Figure S12).

Secondly, we conducted functional connectivity analyses similar to those performed in humans to assess the information transmission between MT/V4 and TEO in monkeys. The DCM analysis showed that the “bidirectional modulation” had the highest posterior probability for both the intact macaque and human stimuli (Figure S14A). Similar to the human results, the driving inputs of the upright BM stimuli were significantly stronger than those of the inverted ones, but only for within-species BM perception (the intact macaque BM stimuli in Figure S14B, *F*_1,51_ = 8.25, *p* = 0.006). This was not the case for cross-species BM perception (the intact human BM stimuli in Figure S14B, *F*_1,56_ < 0.01, *p* = 0.945). However, the modulatory effect of the upright BM was significantly stronger than that of the inverted BM from MT/V4 to TEO for both the intact macaque and human stimuli on the forward connection (Figure S14B, macaque BM: *F*_1,51_ = 21.88, *p* < 0.001; *F*_1,56_ = 8.23, human BM: *p* = 0.006), as well as on the backward connection (Figure S14B, macaque BM: *F_1_*_,51_ = 4.07, *p* = 0.049; human BM: *F*_1,56_ = 5.86, *p* = 0.019). These results suggest that the monkey TEO does not perform the same function as the human pSTS, indicating no functionally homogenous brain clusters in monkey STS are completely comparable to the human pSTS in the current study.

## Discussion

During evolution, animals have developed a remarkable ability to quickly recognize BM both within and across species, which is fundamental to their individual survival and mutual collaboration. The current study investigated the brain mechanisms underpinning species-general and species-specific BM perception. Using a similar experimental design, we recruited both human and monkey participants and searched for the crucial brain areas located in the third visual pathway that selectively processes socially relevant visual cues (Pitcher & Ungerleider, 2021). The results demonstrated that for humans, the hMT+ area responded similarly to both the intact human and macaque BM stimuli, while the pSTS area selectively responded to the intact human BM stimuli. This was further supported by an increased feedforward connection from hMT+ to pSTS only when processing the intact human BM stimuli. For monkeys, the MT area could also indiscriminately represent both the intact human and macaque BM stimuli; however, there were no areas specifically tuned to the macaque BM stimuli. The direct comparison between the findings in humans and monkeys substantiates that the upstream brain region (i.e., MT) on the third visual pathway may develop a generalized ability to encode BM across species, and this ability is functionally homogeneous across species. By contrast, the downstream brain region in STS may show functional divergence across species. The human pSTS may have evolved as a hub for species-specific BM perception. However, the evidence for such BM processing in the monkey STS region was limited. These results revealed shared and unique neural mechanisms underlying BM perception across species during evolution.

Within the third pathway of visual processing, the MT emerges as a primary recipient of visual motion information from the primary visual cortex, prior to the STS during BM perception (Pitcher & Ungerleider, 2021). Previous studies have shown that the hMT+, a region known for its sensitivity to motion, is strongly activated when processing BM information (Grezes et al., 2001). BM stimuli provide a wealth of information, including the shape of the moving subject, the species of the organism, and the intention of the action. Among these, the hMT+ region is predominantly dedicated to deciphering the motion-related aspects (Peelen et al., 2006; Thompson & Baccus, 2012), particularly the complex coherent motion contained in BMs (Peuskens, Vanrie, Verfaillie, & Orban, 2005). Furthermore, the hMT+ can also be activated by scrambled BM or local feet motion, suggesting that the hMT+ is able to process basic motion aspects in the early stages of BM perception, such as acceleration or opponent motion (H. F. Chang et al., 2018; Duarte, Abreu, & Castelo-Branco, 2022; Peuskens et al., 2005). The view that the hMT+ functions as an initial motion analysis of the BM stimuli implicitly implies its general ability to encode the BM cues shared across various species (Duarte et al., 2022; Hirai & Senju, 2020), which has been confirmed by our results.

In the third visual pathway, dynamic visual information is transmitted to the STS via the MT (Pitcher & Ungerleider, 2021). Our results also demonstrated that it is in the pSTS region that the conspecific BM information (i.e., the human BM) is specifically identified. Neurophysiological studies have pinpointed the human pSTS as the region most critically involved in BM perception (for a review, see (Allison, Puce, & McCarthy, 2000); for a meta-analysis, see (Grosbras et al., 2012)). Remarkably, the pSTS is strongly activated when people observe human BMs compared to other motions like scrambled, robot, or object motions (Mar, Kelley, Heatherton, & Macrae, 2007). The species-specific response found in the pSTS is presumably because the conspecific BM inherently carries essential social information for potential communication (Papeo, Wurm, Oosterhof, & Caramazza, 2017). The pSTS has been revealed as a hub that integrates relevant social information and processes a plethora of socially significant stimuli (Lahnakoski et al., 2012), including communicative intention (Redcay, Velnoskey, & Rowe, 2016) and social interaction (Isik et al., 2017). In line with this explanation, preliminary evidence has shown that the pSTS responds more strongly to human motions compared with animal motions (Han et al., 2013; Kaiser, Shiffrar, & Pelphrey, 2012; Papeo et al., 2017). Our results extended the previous findings by applying a stricter contrast (upright BM versus inverted BM) and revealing increased functional connectivity between hMT+ and pSTS during the perception of conspecific BM. The functional connectivity results clearly portrayed that the cross-species BM information initially processed in the hMT+ may be toned down during the transition to the pSTS, particularly due to its limited social relevance for humans. Together, these findings support the notion that the pSTS, as a downstream brain region, may selectively represent the high-level social information contained in within-species BMs.

To trace the evolutionary origins of the neural mechanism underlying BM processing in humans, we conducted a parallel study employing an identical factorial design to investigate brain activity associated with BM perception in macaque monkeys. The existence of the third pathway involved in processing BM has been confirmed in macaques as well (Pitcher & Ungerleider, 2021). Anatomical research has revealed that the monkey MT area, situated on the posterior bank of the superior temporal sulcus, receives direct projection from the primary visual cortex (V1) and connects to the anterior regions of the dorsal bank and fundus of the STS (Boussaoud, Ungerleider, & Desimone, 1990; Ungerleider & Desimone, 1986).

Similar to human studies, the monkey MT area is pivotal for the analysis of visual motion (Maunsell & van Essen, 1983) and can be significantly activated when monkeys watch either human point-light displays (Krekelberg, Augath, & Logothetis, 2002) or monkey movement stimuli (Jastorff et al., 2012). In conjunction with our findings from univariate and multivariate analyses, it is revealed that the macaque MT area is equipped with the ability to detect the typical pendular movement cues present in BM, even from different species. However, the present study did not find any downstream regions in the STS in monkeys that responded more to upright stimuli than inverted ones, either from the same species or different species. On the contrary, previous electrophysiology and imaging studies discovered that the fundus and upper bank of the posterior and middle STS (Jastorff et al., 2012), the mid-anterior STS region (Krekelberg et al., 2002), and even the anterior superior temporal polysensory area (the upper bank of the front part of the STS) (Oram & Perrett, 1994) were involved in processing the kinematics of monkey and human BM stimuli, demonstrating that discontinuous portions of the monkey STS can be activated by both within- and cross-species BM stimuli.

The reason why we did not observe stronger activation to upright BM stimuli relative to inverted BM stimuli in monkey STS may be attributed to the contrast we employed. Different from the contrast between intact and scrambled BM in Krekelberg et al.’s study and the contrast between biological and translational motion in Jastorff et al.’s study, the contrast between upright and inverted BM used in our study selectively captures the kinetics along the direction of gravity (Ma et al., 2022). Unlike humans, monkeys in the natural environment can not only hang on trees in an upside-down position but also accumulate rich visual experience from observing other monkeys hanging upside down. Therefore, monkeys may be equally sensitive to upright and inverted BMs compared to humans. Analogously, humans, if exposed to a microgravity environment for a prolonged period, would increase their sensitivity to inverted BM through decreasing activation in the pSTS (Wang et al., 2022). It is probable that the natural environment to which monkeys are exposed shapes their STS regions’ responses to inverted BMs.

However, are there some subregions in the monkey STS that exhibit species-specific properties when processing BM stimuli? In Jastorff et al.’s investigation, they found the body patches in the STS show stronger activation to conspecifics compared to human action (Jastorff et al., 2012). Although the entire monkey STS region is comparably activated by the upright and inverted versions of BM stimuli in the present study, we wondered whether the activation pattern from a certain proportion of it specifically represents within-species BM rather than cross-species BM. Indeed, we found one region around TEO, which can represent within-species BM but cannot represent the shared BM across species, similar to the human pSTS. However, upon examining the functional connectivity between MT/V4 and TEO, unlike in humans, we observed very similar results for own-species and other-species BM stimuli in monkeys. Hence, the species-specific BM processing (especially with kinetics along the direction of gravity) exists in monkey STS, but it may not employ the same mechanisms as observed in humans. In light of these results, species-specific BM perception in monkeys warrants further investigation.

Previous findings have provided ample evidence that the third visual pathway is dedicated to BM processing in both human and non-human primate brains (Duarte et al., 2022; Jastorff et al., 2012; Pitcher & Ungerleider, 2021). Our study advances beyond previous knowledge by illustrating that the neural codes for BM perception in humans and monkeys are not fully duplicated. Specifically, we identified a neural pathway in humans, from hMT+ to pSTS, that selectively processes conspecific BMs, filtering out BMs from other species. Conversely, in monkeys, BM from all species is generally processed in the MT, without a similar species-specific representation in the STS. The comparison clearly reveals that while the relatively upstream brain regions (i.e., MT) share homologous functions across species, the relatively downstream brain regions in the STS may have undergone differentiation and specialization. This striking difference may reflect distinct evolutionary trajectories of each species after diverging from their common ancestor. Humans, having gradually adapted to living permanently in fixed settlements with their own species, have significantly reduced their interactions with other species. This shift prioritizes their social communication with conspecifics and may have facilitated the divergence of species-specific biosocial information processing. By contrast, monkeys, living in competitive natural environments full of various species, need to be sensitive to all species (e.g., predators) for survival. Consequently, they may not exhibit the same evolution towards species-specific biosocial information processing as observed in humans. Future studies employing more ecological stimuli are needed to corroborate these findings.

## Conclusion

The present study innovatively adopts a cross-species comparison approach to delve into the neural substrates involved in both within-species and cross-species BM perception, as well as its divergence across species. These findings not only enrich our understanding of the evolutionary aspects of the third visual pathway, but also pave the way for future research, particularly in considering species differences in processing biosocial information.

## Method

### Subjects

#### Humans

Twenty-one healthy adults (11 males, mean age 25 ± 3 years) were enrolled in the fMRI experiment. All participants had normal or corrected-to-normal vision and provided written informed consent before the formal experiment. The current study was conducted in accordance with the Declaration of Helsinki and was approved by the institutional review board of the Institute of Psychology, Chinese Academy of Sciences. Two participants were removed because their ROIs (regions-of-interest) could not be reliably found under the predefined threshold, and one participant was removed due to excessive head movements (>2mm). Thus, eighteen participants were included for further analysis.

#### Monkeys

Three male macaque monkeys (monkeys Q, K, and M; Macaca mulatta; 10-11 y old; 7.0 – 10.5 kg) were used. They were acquired from the same primate breeding facility in China, where they had social group histories as well as group-housing experience until their transfer to the Institute of Biophysics (IBP), CAS, for quarantine at the age of approximately 4 y. Animals used in this study had been housed at IBP for 6–7 y before this experiment. All experimental procedures complied with the US National Institutes of Health Guide for the Care and Use of Laboratory Animals and were also approved by the Institutional Animal Care and Use Committee of IBP. Each monkey was surgically implanted with a magnetic resonance (MR)-compatible head post under sterile conditions, using isoflurane anesthesia. After recovery, subjects were trained to sit in a plastic restraint chair and fixate on a central target for long durations with their heads fixed, facing a screen on which visual stimuli were presented (Liu et al., 2022; Liu et al., 2015; Liu et al., 2013).

#### Stimuli

For both the human and monkey fMRI experiments, we used the same set of human and macaque BM stimuli. Human point-light BM stimuli were adopted from motion capture data (Troje, 2002), which consisted of 15 dots located at the major body joints from the head, shoulders, elbows, wrists, hips, knees, and ankles of human actors (see Figure 1A). We then randomly presented six viewpoints of the point-light walkers uniformly distributed from 90° leftwards to 90° rightwards to avoid neural adaptation in the fMRI experiment.

Macaque point-light BM stimuli were converted from three full-body macaque video clips depicting their walking movement from the side. We first manually tracked the motion sequences of 20 major joints in the four legs (3×4), head (3), back (3), and tail (2) of monkeys frame by frame (see Figure 1A). To keep the macaques walking in place, we centralized the positions of the point-light BM stimuli to cancel the translation. Their moving trajectories were further smoothed by realigning the position of each joint at each frame to a virtual position lying linearly between its preceding and following positions, while keeping the distances from this position to its preceding and next positions in the same ratio with the distances calculated using the original positions. This algorithm could effectively eliminate the irregular moving jitters without affecting the core kinetics of biological agents. The three-macaque point-light BM stimuli walked towards the left or right (i.e., six random stimuli comparable to the human BM stimuli).

Additionally, we generated scrambled counterparts for each intact stimulus by randomizing the initial position of each point within the same region. Such manipulation keeps their local motion trajectories intact but disrupts the global configuration. As controls, inverted BM stimuli were created by vertically mirror-flipping the four upright BM sequences (i.e., intact human, scrambled human, intact macaque, and scrambled macaque).

### fMRI acquisition

#### Humans

Functional and anatomical MRI scanning was carried out in the Beijing MRI Center for Brain Research (BMCBR) at a 3-Tesla Siemens Prisma MRI scanner with a 20-channel head coil. For anatomical scans, a high-resolution T1-weighted structural image was obtained with the following parameters: voxel size: 1 mm isotropic; repetition time (TR): 2600 ms; echo time (TE): 3.02 ms; flip angle (FA) = 8°; field of view (FOV): 240 × 240 mm; Functional images were acquired with the following parameters: T2*-weighted gradient - multiband accelerated echo-planar imaging (EPI) sequences; Imaging parameters were as follows: voxel size: 2 mm isotropic TR:2000 ms; TE:30 ms; FA: 70 °; FOV: 192 × 192m.

#### Monkeys

Functional and anatomical MRI scanning was performed with the same equipment as humans with an 8-channel head coil. Before each scanning session, an exogenous contrast agent [monocrystalline iron oxide nano colloid (MION)] was injected into the femoral or external saphenous vein (8 mg/kg) to increase the contrast/noise ratio and to optimize the localization of fMRI signals (Leite et al., 2002). Forty-eight 1.5 mm coronal slices (no gap) were acquired using single-shot interleaved gradient-recalled echo planar imaging. Imaging parameters were as follows: voxel size: 1.5 mm isotropic; TR: 2.5 s; TE: 17 ms; FA: 90°, FOV: 129 × 129 mm. A low-resolution T2 anatomical scan was also acquired in each session to serve as an anatomical reference (0.625 mm × 0.625 mm × 1.5mm; TE: 101 ms; TR: 11.2 s; flip angle: 126°). To facilitate cortical surface alignment and the following local targeting, we also acquired high-resolution T1-weighted whole-brain anatomical scans in separate sessions. Imaging parameters were as follows: voxel size: 0.5 mm isotropic; TR: 2.2 s; TE: 2.84 ms; flip angle: 8°.

### Human fMRI experiments

#### Main experiment

The main fMRI experiment adopted a within-subjects block design. Participants were scanned in four sessions: scrambled human BM, intact human BM, scrambled macaque BM, and intact macaque BM. The session order was counterbalanced across participants, but scans of scrambled stimuli were always before scans of intact stimuli to avoid recognizing scrambled stimuli once intact stimuli were shown. Each session had two runs. In each run, eight blocks of upright and inverted BM displays were presented in a counterbalanced order. Each block lasted 12s, followed by a 6s interval. During each block, taking intact human BM as an example, participants viewed six exemplars of random BM stimuli from different viewpoints, which were consecutively presented for 2s each. A red fixation (0.5° × 0.5°) appeared in the center against the gray screen during the whole run. Human point-light BM and macaque point-light BM stimuli are subtended approximately 2.3°× 6.6°and 5.3° × 3.6° in visual angle, respectively. To further avoid neural adaptation, we allowed the BM stimuli to float with a random offset of fixation within an area of 1°. Participants were required to maintain their attention on the central fixation and to press a response button when the fixation changed its color to a light red. In the scanner, all stimuli were presented using MATLAB (MathWorks, Inc.) with the Psychophysics toolbox extensions (Brainard & Vision, 1997). Participants viewed the screen (60 Hz frame rate) through a mirror mounted on the head coil.

#### Localizer

To eliminate data circularity issues (Kriegeskorte, Simmons, Bellgowan, & Baker, 2009), participants completed three additional localizer scans after the main experiment. Two functional localizer tasks were performed to define the human and macaque BM-sensitive STS regions, respectively, by contrasting the upright intact human/macaque BM with its inverted scrambled counterpart. In each task, there were two runs, each containing 8 upright intact BM blocks and 8 inverted scrambled BM blocks. All the other procedures of these tasks were identical to the main experiment. The third localizer task was used to define the hMT+. The design of this task was also identical to the main experiment except that the BM stimuli were replaced by moving optic flow (expanding and contracting) and static dots, which were presented within a circular region of radius 3.2° (Huk et al., 2002). Stimuli were shown in 12 blocks in each run (each condition contained 6 blocks) in a counterbalanced order, with a total of two runs.

### Monkey fMRI experiments

#### Main experiment

The main fMRI experiment shared almost the same design and procedure as the human experiment. Here, we only briefly introduced the critical and dissimilar parts. As human participants, three monkeys also engaged in four types of BM stimuli: scrambled human BM, intact human BM, scrambled macaque BM, and intact macaque BM. In each session, 14–21 runs were collected from each animal, and the detailed information was presented in Table S7. In each run, the upright and inverted BM stimuli were displayed in 4 blocks, respectively, in a counterbalanced order. Each block had a duration of 30s, followed by a 15s fixation interval. Six BM stimuli with different viewpoints were consecutively presented for 2.5s and repeated twice during each block. The human BM and macaque BM stimuli are subtended approximately 3.6°×10.3° and 9.1° × 6.2° in visual angle, respectively.

Monkeys were required to maintain fixation on a central square on the screen during the whole experiment and receive a liquid reward. The monkeys’ eye position was tracked using an infrared pupil tracking system (ISCAN, Inc). The frequency of reward would be increased as the duration of fixation increased (Liu et al., 2022; Liu et al., 2015; Liu et al., 2013). The fMRI data enrolled in subsequent analysis only came from those blocks/runs with qualified fixation percentage (Monkey Q, upright: 89.90 ± 0.58%, inverted: 89.74 ± 0.62%; Monkey K, upright: 89.00 ± 0.57%, inverted: 88.38 ± 0.61%; Monkey M, upright: 90.54 ± 0.58%, inverted: 90.69 ± 0.62%.

#### Localizer

Monkeys performed three localizer tasks as humans. The BM stimuli used to define the ROIs sensitive to human and macaque BM stimuli and the moving dots used to define the ROI sensitive to pure physical motion were identical to the localizer tasks of humans. All other parameters were the same as those in the main fMRI experiment of monkeys. The run numbers collected in the three localizers tasks were presented in Table S7 as well.

### Human Data analysis

#### Preprocessing

Preprocessing was implemented with SPM12 (http://www.fil.ion.ucl.ac.uk/spm), and ROI analysis was performed with MarsBar (http://marsbar.sourceforge.net). The imaging data preprocessing involved re-slicing, co-registration, segmentation, normalization, and smoothing. Specifically, EPI volumes were realigned using the first volume of each run as a reference. Then, the structural image was coregistered to the reference image and segmented into gray matter, white matter, and CSF in each individual space. The segment parameters were further used to normalize each individual’s functional images onto MNI space. After normalization, data were spatially smoothed with an FWHM, 4 mm Gaussian kernel.

#### Whole-brain analysis

All data were fitted through a general linear model (GLM). For each type of BM stimuli (i.e., scrambled human, intact human, scrambled macaque, intact macaque), the GLM analyses included two regressors of interest (upright and inverted versions) and six motion regressors of no interest (three translation parameters and three rotation parameters). All regressors were convolved with the hemodynamic response function. Two contrasts, upright > inverted and inverted > upright, were computed for each participant in a first-level t-test. To show group-level statistical maps across subjects, contrast images of all the participants were included in a second-level group analysis, with an FDR-corrected threshold of *p* < 0.05 at the cluster level (*p* < 0.001 uncorrected at the voxel level) and an extent threshold of 10 voxels.

#### ROI analysis

We identified the bilateral pSTS regions sensitive to human BM with the contrast of upright intact human BM > inverted scrambled human BM (*p* < 0.05, uncorrected). Based on both the anatomical template and statistics maps, each individual ROI was defined as a 6 mm radius sphere centered on the individual local maxima within the designated region (i.e., superior temporal cortex). In the same vein, we identified the bilateral pSTS regions sensitive to macaque BM with the contrast of upright intact macaque BM > inverted scrambled macaque BM (*p* < 0.05, uncorrected) and the hMT+ with the contrast of moving dots > static dots (*p* < 0.005, uncorrected). We extracted the beta values of the BOLD signal within all the ROIs for each participant. These values were further analyzed by a 2 (hemisphere: left and right) × 2 (orientation: upright and inverted) repeated-measures ANOVA for each type of BM stimuli. The ROIs were also applied in the subsequent PPI and DCM analysis. Their mean MNI coordinates for all participants were described in Supplementary Table 1, and the locations of these ROIs for each participant are displayed in Supplementary Materials (Figure S1).

#### MVPA analysis

To further examine which brain regions encode the upright and inverted BM stimuli, we conducted a multivariate pattern analysis (MVPA) implemented in the COSMOMVPA toolbox (Oosterhof, Connolly, & Haxby, 2016). In this analysis, the functional images had not been normalized and spatially smoothed with a Gaussian kernel of 2 mm. ROIs were created in individual native spaces. We trained the support vector machine (SVM) classifier (C.-C. Chang & Lin, 2011) to distinguish between upright and inverted BM stimuli in a block-based fashion. We trained the classifier on data from 31 out of 32 blocks (leaving one out). We assessed the accuracy of the classifier by testing it on the upright/inverted BM stimuli in the held- out block. That was iterated 32 times, and the classification accuracy was averaged for each subject and ROI. These accuracies were then entered into one-sample t-tests and were compared to a 50% chance level (α = 0.05, two-tailed, Bonferroni corrected). The above analysis was separately conducted for scrambled human, intact human, intact macaque, and scrambled macaque BM stimuli.

To assess whether the representation of BM is generalized across species, we conducted a cross-classification MVPA. The classifier was trained to discriminate the upright and inverted version of the intact human BM stimuli but tested on its accuracy at classifying the upright and inverted version of the intact macaque BM stimuli and vice versa. The same approach was repeated for the scrambled BM stimuli. For each ROI, individual classification accuracies were subsequently compared against the chance level with one-sample t-tests (α = 0.05, two-tailed, Bonferroni corrected). Significant cross-classification accuracy demonstrates shared representation underlying human and macaque BM perception.

#### PPI and DCM analysis

To investigate the functional coupling among brain regions related to BM perception, we conducted an exploratory PPI analysis (Friston et al., 1997) and selected the bilateral hMT+ as the seed region. We selected the previously defined bilateral hMT+ as a seed region and extracted the first eigenvariate time series from them as the physiological term. The interaction term of PPI analysis was calculated by multiplying the hMT+ activity and psychological variable of interest (upright vs. inverted) (Gitelman, Penny, Ashburner, & Friston, 2003). Afterward, the convolved regressor for the interaction term, the physiological variable (i.e., the hMT+ activity), and the psychological variable were entered into a first-level GLM analysis, where the six head movement parameters were treated as nuisance regressors. The interaction terms for individuals’ contrast images were further estimated to create group-level maps (*p* < 0.05 after FWE correction).

Given that PPI analysis does not reveal the direction of the functional connectivity between brain regions, we conducted DCM analysis in the DCM tool of SPM12. Based on previous studies and the key implication of our PPI results, we focused on the effective connectivity between the hMT+ and pSTS regions when intact human and macaque BMs were displayed. It is worth noting that in the DCM analysis, the human pSTS ROI was used for intact human BM display, while the macaque pSTS ROI was used for intact macaque BM display. For each participant, the same ROIs in the left and right hemispheres were combined for further analysis. Then, we extracted the activity time series for each volume of interest by computing the first eigenvector of all its voxels.

We constructed the DCM models by first assuming that the driving visual input (here, both the upright and inverted BM stimuli) through hMT+ entered the system (C-matrix). Second, we assumed the intrinsic connection between the hMT+ and the pSTS was bidirectional (hMT+ ↔ pSTS; A-matrix). To reveal how BM perception modulates the endogenous connectivity between hMT+ and pSTS, we systematically varied the modulatory effects of BM perception (upright versus inverted) on the intrinsic connections between MT and pSTS (B-matrix) in 4 models: (1) the “no modulation” mode; (2) the “forward modulation” mode, in which the connection from hMT+ to pSTS is unidirectionally modulated by BM perception; (3) the “backward modulation” mode, in which the connection from pSTS to hMT+ is unidirectionally modulated by BM perception; (4) the “bidirectional modulation” mode, in which both feedforward and feedback connections between hMT+ and pSTS are modulated by BM perception. For illustrative purposes, all the hypothesized models were drawn in Figure 4A. Then, we conducted the model comparison using a fixed-effects Bayesian model selection (BMS), and the most appropriate model was assessed by exceedance probability (Stephan, Penny, Daunizeau, Moran, & Friston, 2009). The parameter values of the winning model relating to the modulation effect were assessed using paired t-tests at the group level.

### Monkey Data analysis

#### Preprocessing

Image preprocessing was conducted with AFNI (Cox, 1996). Briefly, the EPI volumes were time-corrected and realigned to the base volume with minimum outliers. Then, outliers of EPI images were eliminated with the 3dDespike function. EPI volumes were next smoothed with a 2mm-full-width half-maximum Gaussian kernel and normalized to the mean signal value within each run. Finally, EPI volumes were coregistered with the T1 volume to the NIMH Macaque Template (NMT v2) space (Jung et al., 2021).

#### Whole-brain analysis

After preprocessing, a GLM was fitted to estimate and compare the neural activities between the upright and inverted versions for each type of BM stimuli (i.e., scrambled/intact human, scrambled/intact macaque). In addition to the above regressor of interest, each GLM model included a hemodynamic response function with the MION response curve and regressors with no interests (baseline, movement parameters from realignment corrections, and signal drifts). For each type of BM stimuli, the contrast map (upright vs. inverted) was computed for each monkey with an FDR-corrected threshold of *ps* < 0.05 at the cluster level (*p* < 0.001 uncorrected at the voxel level and an extent threshold of 5 voxels). Finally, a cluster would be determined as significant if it was found in both the left and right hemispheres, as well as across all three monkeys. All fMRI signals throughout the study were inverted so that an increase in signal intensity indicates an increase in activation (Vanduffel et al., 2001).

#### ROI analysis

The monkey ROI analysis was performed using similar contrasts as humans (e.g., upright intact BM > inverted scrambled BM or moving dots > static dots) and adjusted for the following reasons. First, as almost all areas along the monkey STS exhibited significant differences in the MT functional localizer (FDR corrected *p* < 0.05), we selected the top 60 voxels that fell within the MT and V4t anatomical area (dilated by one voxel using AFNI 3dmask_tool) at each hemisphere by using the D99 macaque brain atlas (Reveley et al., 2017) and named the cluster the MT/V4 as a comparable ROI to human hMT+ (Jastorff et al., 2012; H. Kolster et al., 2009; Hauke Kolster, Peeters, & Orban, 2010; Ungerleider & Desimone, 1986). Second, in both the human and monkey BM localizers, we were not able to find a functionally homogenous cluster in the monkeys comparable to the pSTS in humans, which means no clusters in the STS responded more strongly to the upright intact BM stimuli than the inverted scrambled BM stimuli (except those areas close to and overlapped with the MT/V4 ROI, see Figure S3). Therefore, only the MT/V4 ROI was localized for the monkeys, with detailed information in Table S8 and Figure S4. To compare the beta values of the MION signals from the upright and inverted version, a 2 (hemisphere: left; right) × 2 (orientation: upright; inverted) LMM with runs, sessions, and monkeys as random factors was run on the MT/V4 ROI (Liu et al., 2022) for each type of BM stimuli respectively (i.e., scrambled/intact human, scrambled/intact macaque). All the models were fitted in the R programming version 3.6.3.

#### MVPA analysis

The MVPA was also conducted using the COSMOMVPA toolbox. For each monkey, we trained an SVM classifier using data from the predefined ROI to distinguish between the upright and inverted versions for each type of BM stimuli in a run-based fashion. The leave-one-run-out cross-validation was used to get the classification accuracy (Dubois, de Berker, & Tsao, 2015), and this process was iterated N times (N was the total number of runs for each monkey). Following the human analysis, such manipulation was conducted within and across species. Due to the small sample size in the monkey experiments, we used a binomial test to evaluate the statistical significance of the classification accuracy (Dubois et al., 2015) and enrolled all runs from 3 monkeys for the group analysis. All these statistical tests were performed in the R programming version 3.6.3.

## Acknowledgements

We are grateful to Professor Nikolaus Troje for kindly providing us with the human BM stimuli and Dr. Zhaoqi Hu for his assistance and suggestions in experimental design. This research was supported by grants from the STI2030-Major Projects (2021ZD0203800, 2021ZD0200200), the National Natural Science Foundation of China (31830037, 31600884), the Interdisciplinary Innovation Team (JCTD-2021-06), Youth Innovation Promotion Association of the Chinese Academy of Sciences, and the Fundamental Research Funds for the Central Universities.

## Data and code availability

All data and code that supports the findings of this study are available from Psychological Science Bank (https://www.scidb.cn/en/s/RZz6vu).

