## Supplementary materials for "Shared and Unique Neural Codes for Biological Motion Perception in Humans and Macaque Monkeys"

### Supplementary material

#### Section A: Supplemental fMRI results

##### *Human results*

**Table S1.** Clusters showing stronger activation in upright > inverted or inverted > upright contrasts in the four types of BM stimuli. Peak coordinates are in the Montreal Neurological Institute space (MNI). For each cluster, anatomical labels, MNI coordinates, z score, cluster size in the number of voxels, and *p* value are displayed ($p < 0.001$, cluster threshold of $p < 0.05$ (FWE), and the minimum cluster size > 10 voxels).

| Regions | MNI coordinates peak | | | Z score | Cluster Size (voxels) | *p* value (FWE-corr) |
|---|---|---|---|---|---|---|
| | X | Y | Z | | | |
| **Human BM(Intact)** | | | | | | |
| ***Upr>Inv*** | | | | | | |
| Left Inferior Frontal Gyrus | 46 | 4 | 30 | 4.85 | 67 | **0.001** |
| Right Postcentral Gyrus | 16 | -40 | 66 | 4.6 | 46 | **0.013** |
| Right Middle Temporal Gyrus | 50 | -64 | 10 | 4.49 | 463 | **0.001** |
| Right Lingual Gyrus | 18 | -84 | -6 | 4.16 | 122 | **0.001** |
| Left Lingual Gyrus | -20 | -88 | -10 | 4.14 | 111 | **0.001** |
| Right Fusiform Gyrus | 38 | -50 | -22 | 4.09 | 30 | 0.1 |
| Right Cuneus | 12 | -88 | 32 | 4.05 | 72 | **0.001** |
| Left Precuneus | -18 | -62 | 46 | 4 | 30 | 0.1 |
| Left Middle Occipital Gyrus | -34 | -74 | 0 | 3.95 | 119 | **0.001** |
| Left Inferior Parietal Lobule | -38 | -42 | 48 | 3.76 | 34 | 0.059 |
| ***Inv>Upr*** | | | | | | |
| No suprathreshold clusters | | | | | | |
| **Macaque BM(Intact)** | | | | | | |
| ***Upr>Inv*** | | | | | | |
| Right Cuneus | 14 | -90 | 26 | 5.26 | 169 | **0.001** |
| Left Cuneus | -10 | -94 | 22 | 4.52 | 88 | **0.001** |
| Middle Occipital Gyrus | 46 | -78 | 4 | 4.26 | 62 | **0.006** |
| ***Inv>Upr*** | | | | | | |
| Lingual Gyrus | -2 | -80 | 0 | 5.44 | 492 | **0.001** |

| | | | | | | |
|---|---|---|---|---|---|---|
| **Human BM(Scrambled)** | | | | | | |
| ***Upr>Inv*** | | | | | | |
| No suprathreshold clusters | | | | | | |
| ***Inv> Upr*** | | | | | | |
| No suprathreshold clusters | | | | | | |
| **Macaque BM(Scrambled)** | | | | | | |
| ***Upr>Inv*** | | | | | | |
| Left Cuneus | -16 | -92 | 20 | 5 | 129 | **0.001** |
| Left Cerebelum_Crus1_L (aal) | -30 | -86 | -24 | 4.56 | 38 | **0.035** |
| Left Superior Frontal Gyrus | -36 | 22 | 56 | 4.14 | 41 | **0.024** |
| Left Supramarginal Gyrus | -58 | -56 | 28 | 4.08 | 36 | **0.046** |
| Right Cuneus | 12 | -94 | 16 | 3.95 | 95 | **0.001** |
| ***Inv>Upr*** | | | | | | |
| Lingual Gyrus | -8 | -80 | -6 0 | 4.54 | 571 | **0.001** |

**Table S2.** Mean MNI coordinates for the defined ROIs across all participants

| ROI | X | Y | Z |
|---|---|---|---|
| ***Right hemisphere*** | | | |
| **Human-pSTS** | 54 | -51 | 15 |
| **Macaque-pSTS** | 54 | -49 | 15 |
| **hMT+** | 47 | -68 | 8 |
| ***Left hemisphere*** | | | |
| **Human-pSTS** | -52 | -48 | 13 |
| **Macaque-pSTS** | -52 | -49 | 14 |
| **hMT+** | -45 | -70 | 8 |

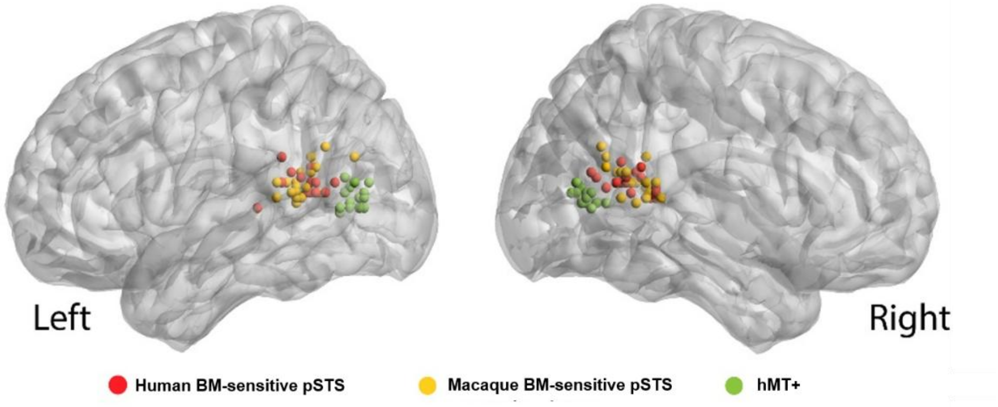

**Figure S1**. Locations of the predefined ROIs for each participant on a standard brain (MNI template). The human BM-sensitive pSTS are shown in red, the macaque BM-sensitive pSTS is shown in yellow, and the hMT+ is shown in green.

**Table S3.** Statistical results of main and interaction effects of ANOVAs on the mean activation in the hMT+ and pSTS ROIs. $p < 0.05$ in bold.

| | hMT+ | | | | | | pSTS | | | | | |
|---|---|---|---|---|---|---|---|---|---|---|---|---|
| | Interaction | | orientation | | hemisphere | | interaction | | orientation | | hemisphere | |
| | F | *p* | F | *p* | F | *p* | F | *p* | F | *p* | F | *p* |
| **Intact human BM** | 9.18 | **0.008** | 6.16 | **0.024** | 0.02 | 0.878 | 6.36 | **0.022** | 18.25 | **<0.001** | 2.72 | **0.118** |
| **Intact macaque BM** | 11.61 | **0.003** | 3.91 | 0.064 | 0.04 | 0.846 | 0.08 | 0.782 | 0.02 | 0.879 | 0.1 | **0.750** |
| **Scrambled human BM** | 0.24 | 0.632 | 0.95 | 0.343 | 0 | 0.966 | 0.8 | 0.384 | 2.29 | 0.149 | 0.78 | **0.390** |
| **Scrambled macaque BM** | 7.47 | **0.01** | 2.48 | 0.134 | 0.28 | 0.603 | 0.41 | 0.528 | 5.69 | **0.03** | 0.38 | **0.546** |

**Table S4.** Statistical results of within-species MVPA in the hMT+ and pSTS ROIs. $p < 0.05$ in bold.

| | hMT+ | | | | | | pSTS | | | | | |
|---|---|---|---|---|---|---|---|---|---|---|---|---|
| | Left | | | Right | | | Left | | | Right | | |
| | ACC | T | *p* | ACC | T | *p* | ACC | T | *p* | ACC | T | *p* |
| **Intact human BM** | 66.15 | 5.78 | **<0.001** | 71.5 | 7.59 | **<0.001** | 52.4 | 1.06 | >0.9 | 58.3 | 2.84 | **0.045** |
| **Intact macaque BM** | 69.3 | 8.51 | **<0.001** | 71.2 | 6.83 | **<0.001** | 61.1 | 3.91 | **0.005** | 54.7 | 1.79 | 0.364 |
| **Scrambled human BM** | 53.3 | 1.51 | 0.595 | 59.6 | 3.97 | **0.004** | 53.3 | 1.30 | 0.849 | 54.5 | 2.92 | **0.038** |
| **Scrambled macaque BM** | 52.4 | 0.90 | >0.9 | 55.2 | 2.01 | 0.241 | 0.55 | 2.17 | 0.179 | 55.9 | 3.61 | **0.009** |

**Table S5.** Statistical results of cross-species MVPA in the hMT+ and pSTS ROIs. $p < 0.05$ in bold.

| | | hMT+ | | | | | | pSTS | | | | | |
|---|---|---|---|---|---|---|---|---|---|---|---|---|---|
| | | Left | | | Right | | | Left | | | Right | | |
| **Train** | **Test** | ACC | T | *p* | ACC | T | *p* | ACC | T | *p* | ACC | T | *p* |
| **Intact human BM** | **Intact macaque BM** | 54.0 | 2.31 | 0.135 | 57.5 | 3.24 | **0.019** | 49.1 | -0.70 | >0.9 | 50.9 | 55.2 | >0.9 |
| **Intact macaque BM** | **Intact human BM** | 52.4 | 1.61 | 0.500 | 55.2 | 2.86 | **0.044** | 49.1 | -0.75 | >0.9 | 53.0 | 1.48 | 0.631 |
| **Scrambled human BM** | **Scrambled macaque BM** | 50.5 | 0.47 | >0.9 | 49.7 | -0.17 | >0.9 | 47.2 | -1.70 | 0.430 | 49.8 | -0.17 | >0.9 |
| **Scrambled macaque BM** | **Scrambled macaque BM** | 50.0 | 0 | >0.9 | 50.9 | 0.57 | >0.9 | 50.0 | 0 | >0.9 | 50.2 | 0.80 | >0.9 |

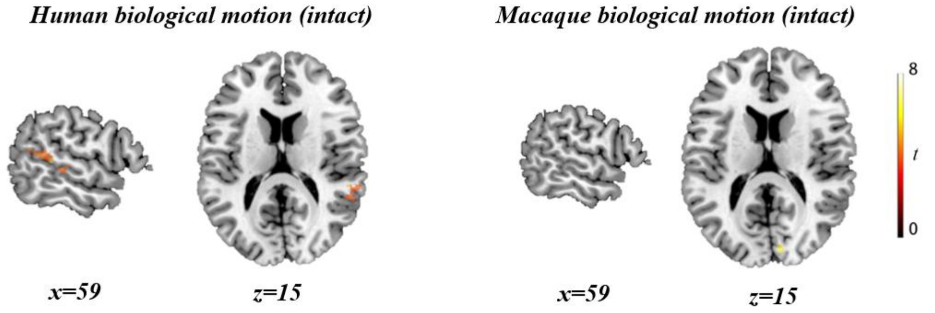

**Figure S2**. Group-level PPI maps of intact human BM with an FDR-corrected threshold of $p < 0.05$ at the cluster level and $p < 0.001$ uncorrected at the voxel level (the minimum cluster size > 10 voxels).

**Table S6.** Coordinates for brain areas showing functional connectivity with the bilateral hMT+ for intact human and macaque BM perception.

| Regions | MNI coordinates peak | | | Z score | Cluster size (voxels) | *p* value |
|---|---|---|---|---|---|---|
| | X | Y | Z | | | |
| **Human BM(Intact)** | | | | | | |
| Right Superior Temporal Gyrus | 64 | -36 | 6 | 5.16 | 142 | **0.001** |
| Left Superior Frontal Gyrus | -4 | 38 | 58 | 4.42 | 110 | **0.001** |
| Right Cingulate Gyrus | 8 | 0 | 36 | 4.16 | 46 | **0.007** |
| Left Cingulate Gyrus | -8 | -40 | 44 | 3.81 | 37 | **0.023** |
| Left Anterior Cingulate | -2 | 48 | 6 | 4.01 | 72 | **0.001** |
| Left Supramarginal Gyrus | -44 | -52 | 28 | 3.84 | 48 | **0.005** |
| Right Middle Temporal Gyrus | 56 | -30 | 0 | 3.84 | 38 | |
| **Macaque BM(Intact)** | | | | | | |
| No suprathreshold clusters | | | | | | |

##### *Monkey results*

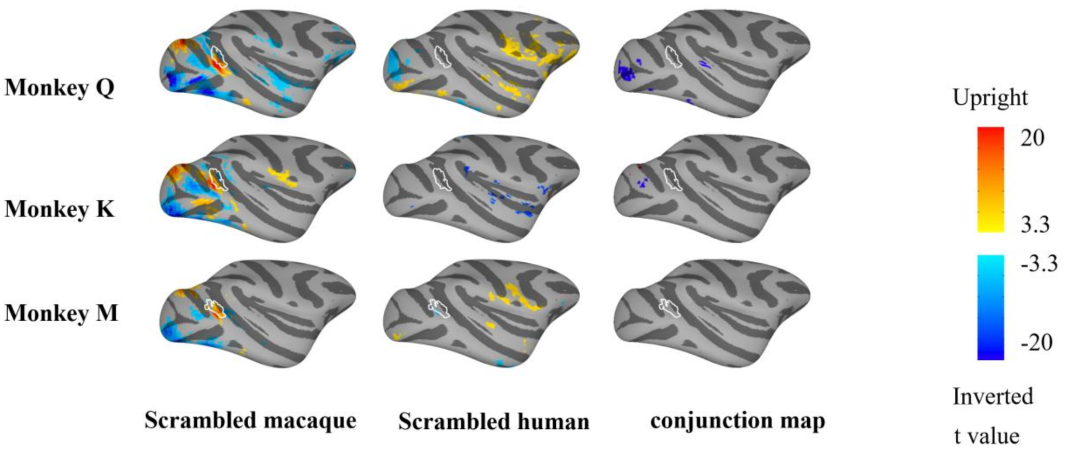

**Figure S3**. Brain activation maps of three monkeys (upright versus inverted) for the scrambled BM stimuli, shown on lateral views of the inflated cortex of the right hemisphere. Panels from top to down represent the results of monkeys Q, K, and M, respectively. Panels from left to right show results from the scrambled macaque and human (uncorrected $p$ < 0.001, FDR corrected $p$ < 0.05), and conjunction maps based on them (FDR corrected $p$ < 0.05), respectively. The borders of the MT/V4 ROI are encircled by white lines.

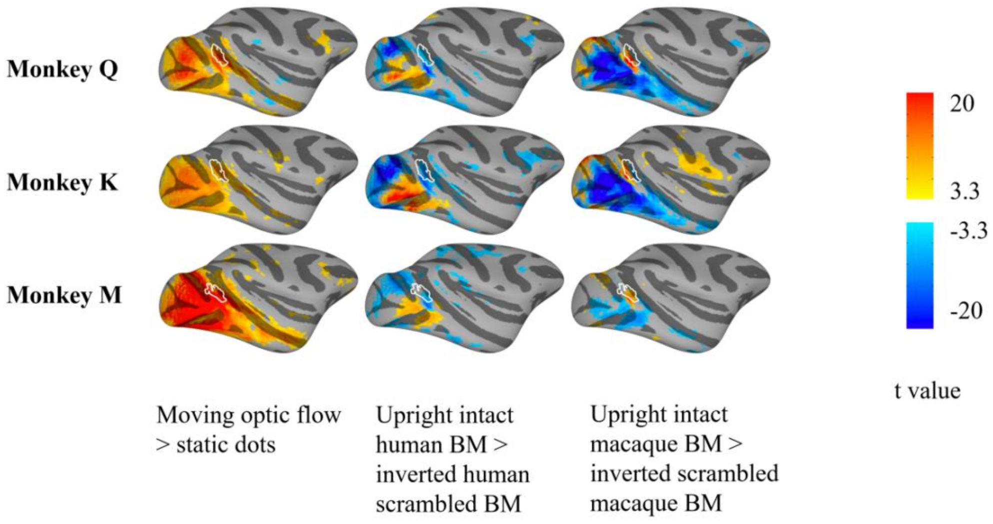

**Figure S4.** Activation maps of three monkeys in the localizer experiments. Panels from top to down represent the results of monkeys Q, K, and M, respectively. Panels from left to right show results from the MT localizer, BM localizer with human stimuli, and BM localizer with macaque stimuli, respectively. uncorrected $p$ <0.001, FDR corrected $p$ <0.05. The borders of the MT/V4 are encircled by white lines.

**Table S7.** Information of the number of sessions and runs conducted in each monkey.

| | Monkey Q | | Monkey K | | Monkey M | |
|---|---|---|---|---|---|---|
| **Number** | Session | Run | Session | Run | Session | Run |
| **Main Experiments** | | | | | | |
| Macaque BM (Intact) | 3 | 18 | 4 | 18 | 2 | 16 |
| Human BM (Intact) | 3 | 18 | 4 | 21 | 3 | 18 |
| Macaque BM (Scrambled) | 3 | 16 | 4 | 18 | 3 | 15 |
| Human BM (Scrambled) | 3 | 14 | 3 | 17 | 3 | 16 |
| **Localizer Experiments** | | | | | | |
| MT | 2 | 8 | 2 | 7 | 2 | 8 |
| Macaque-STS | 1 | 14 | 1 | 14 | 2 | 14 |
| Human-STS | 2 | 13 | 2 | 16 | 2 | 14 |

**Table S8.** NMT coordinates for defined ROIs in each monkey

| ROI | Monkey | X | Y | Z |
|---|---|---|---|---|
| ***Right hemisphere*** | | | | |
| **MT/V4** | | | | |
| | Q | -20.13 | 3.50 | 20.25 |
| | K | -19.38 | 4.25 | 20.50 |
| | M | -21.63 | 4.25 | 21.25 |
| **TEO** | | | | |
| | Q | -25.50 | 0.13 | 13.88 |
| | K | -25.50 | -1.38 | 16.88 |
| | M | -24.63 | -0.75 | 13.50 |
| ***Left hemisphere*** | | | | |
| **MT/V4** | | | | |
| | Q | 20.88 | 4.00 | 20.25 |
| | K | 19.88 | 3.75 | 20.25 |
| | M | 20.13 | 4.50 | 20.75 |
| **TEO** | | | | |
| | Q | 25.50 | 0.13 | 15.38 |
| | K | 25.50 | 0.13 | 15.38 |
| | M | 25.63 | -0.50 | 11.00 |

**Table S9.** Statistical results of main and interaction effects of LMMs on the mean activation in the MT/V4 and TEO ROIs. $p < 0.05$ in bold.

| | MT/V4 | | | | | | TEO | | | | | |
|---|---|---|---|---|---|---|---|---|---|---|---|---|
| | interaction | | orientation | | hemisphere | | interaction | | orientation | | hemisphere | |
| | F | *p* | F | *p* | F | *p* | F | *p* | F | *p* | F | *p* |
| Intact human BM | 0.02 | 0.891 | 11.16 | **0.001** | 0.036 | 0.850 | 0.65 | 0.422 | <0.01 | 0.996 | 0.02 | 0.894 |
| Intact macaque BM | 0.06 | 0.808 | 59.08 | **<0.001** | 1.27 | 0.261 | 0.21 | 0.651 | 0.68 | 0.411 | 0.09 | 0.767 |
| Scrambled human BM | 0.21 | 0.645 | 0.20 | 0.659 | 1.06 | 0.304 | 0.17 | 0.681 | <0.01 | 0.953 | 4.74 | **0.031** |
| Scrambled macaque BM | 0.09 | 0.761 | 57.60 | **<0.001** | 30.44 | **<0.001** | 0.02 | 0.885 | 0.46 | 0.478 | 9.89 | **0.002** |

**Table S10.** Statistical results of within-species MVPA in the MT/V4 and TEO ROIs. $p < 0.05$ in bold.

| | MT/V4 | | | | TEO | | | |
|---|---|---|---|---|---|---|---|---|
| | Left | | Right | | Left | | Right | |
| | ACC | *p* | ACC | *p* | ACC | *p* | ACC | *p* |
| Intact human BM | 82.5 | **<0.001** | 75.4 | **<0.001** | 76.3 | **<0.001** | 65.8 | **<0.001** |
| Intact macaque BM | 95.2 | **<0.001** | 96.2 | **<0.001** | 74.0 | **<0.001** | 73.1 | **<0.001** |
| Scrambled human BM | 53.2 | 0.606 | 44.7 | >0.9 | 47.9 | >0.9 | 50.0 | >0.9 |
| Scrambled macaque BM | 93.9 | **<0.001** | 95.9 | **<0.001** | 73.5 | **<0.001** | 69.4 | **<0.001** |

**Table S11.** Statistical results of cross-species MVPA in the MT/V4 and TEO ROIs. $p < 0.05$ in bold.

| | | MT/V4 | | | | TEO | | | |
|---|---|---|---|---|---|---|---|---|---|
| | | Left | | Right | | Left | | Right | |
| Train | **Test** | ACC | *p* | ACC | *p* | ACC | *p* | ACC | *p* |
| Intact human BM | **Intact macaque BM** | 74.0 | **<0.001** | 81.7 | **<0.001** | 49.0 | >0.9 | 43.3 | >0.9 |
| Intact macaque BM | **Intact human BM** | 64.0 | **0.004** | 71.9 | **<0.001** | 47.4 | >0.9 | 45.6 | >0.9 |
| Scrambled human BM | **Scrambled macaque BM** | 58.2 | 0.129 | 44.9 | >0.9 | 54.1 | 0.480 | 52.0 | 0.762 |
| Scrambled macaque BM | **Scrambled human BM** | 60.6 | 0.050 | 51.1 | >0.9 | 54.3 | 0.471 | 45.7 | >0.9 |

#### Section B: Results of scrambled BM stimuli

##### *Human results*

###### *Univariate Results*

The whole-brain analysis showed that the comparison between upright and inverted BM did not reveal any significantly increased brain activation for the scrambled human BM stimuli. However, there was notably stronger activation observed in the lateral occipital cortex when participants viewed scrambled macaque stimuli (Table S1).

Next, we performed 2 Hemisphere (left vs. right) × 2 Orientation (upright vs. inverted) repeated measures ANOVAs. Consistent with the whole-brain results, we found no significant effects in the hMT+ and pSTS for the scrambled human BM stimuli (see Figure S5 and Table S3, $Fs < 0.95$, $ps > 0.343$). For the scrambled macaque BM stimuli, there was a significant interaction in the hMT+ ($F_{1,17} = 7.47$, $p = 0.014$). The right hMT+ showed a significant activation for upright BM ($t_{17} = 2.18$, $p = 0.043$). Notably, although the effect of orientation in the pSTS reached significance ($F_{1,17} = 5.69$, $p = 0.029$), it was significantly deactivated and not a concern for the current study.

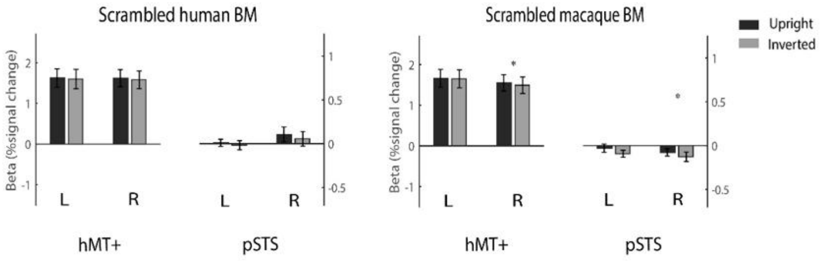

**Figure S5.** The human ROI results. Mean beta values across participants, in each individually defined left and right hMT+ and pSTS, in response to upright versus inverted BM, respectively. Error bars denote standard errors of the mean. *$p < 0.05$.

###### *Multivariate Results*

For within-species decoding results, we found that for the scrambled human BM, classification accuracy was significantly higher than chance level in the right hMT+ (59.5 ± 10.2%, $t_{(17)} = 3.97$, $p = 0.004$) and the right pSTS (54.5 ± 6.5%, $t_{(17)} = 2.93$, $p = 0.037$), while for the scrambled macaque BM, the classification accuracy was only above chance level in the right pSTS (55.9 ± 6.9%, $t_{(17)} = 3.61$, $p = 0.009$; see Figure S6 and Table S4). For cross-species decoding results, the classification accuracies were not significantly higher than chance level for the scrambled human/macaque BM in either the hMT+ or pSTS (*ts* < 0.58, *ps* > 0.9).

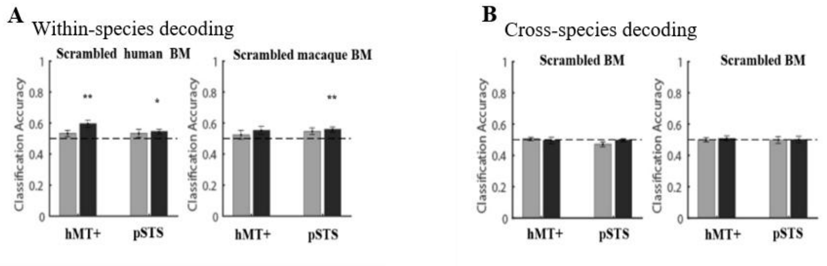

**Figure S6.** The human MVPA results. The classification accuracies of within-species (A) and cross-species (B) decoding of upright and inverted BM stimuli in the hMT+ and pSTS. The horizontal dashed line represents the chance level (50%). Error bars denote standard errors of the mean. *$p < 0.05$, **$p < 0.01$.

#### *Monkey results*

##### *Univariate Results*

For the scrambled macaque BM stimuli, the MT, V4t, and V4 areas also showed increased neural activities for the upright stimuli compared to the inverted stimuli across the three monkeys, whereas for the scrambled human BM stimuli, no consistent brain regions showed significantly stronger activations for the upright stimuli compared to the inverted ones (Figure S3). The ROI results also showed significant main effects of Orientation for the scrambled macaque BM stimuli ($F_{1,144}$= 57.60, *ps* < 0.001), with stronger activation elicited by the upright BM stimuli compared to the inverted ones in the bilateral MT/V4 (left: $t_{144}$ = 5.58, *ps* < 0.001; right: $t_{144}$ = 5.15, *ps* < 0.001). However, there was no significant main effect of Orientation for the scrambled human BM stimuli (see Figure S7, $F_{1,138}$ = 0.20, $p$ = 0.659).

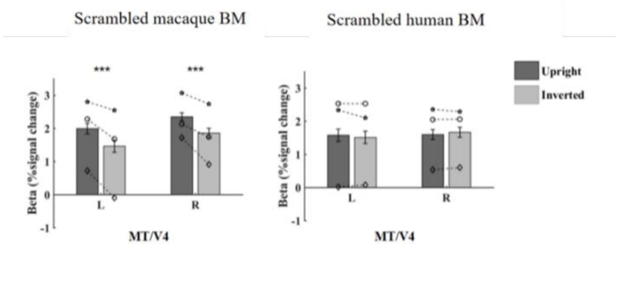

**Figure S7.** The monkey ROI results. Results of the ROI analysis in the MT/V4. Error bars indicate standard errors of the mean. The symbols (circle, star, and diamond) within individual bars represent results from monkeys Q, K, and M, respectively. ***, $p < 0.001$.

##### *Multivariate Results*

For within-species decoding results, the classification accuracies in the bilateral MT/V4 were significantly higher than chance level for the scrambled macaque BM, but not for the scrambled human BM stimuli (see Table S10 for the precise statistical values). Furthermore, we performed cross-species MVPA, only the decoding accuracy of the left MT/V4 reached marginal significance for scrambled BM (by training on the macaque BM and testing on the human BM, left: 60.64 ± 5.04%, $p = 0.050$) (see Figure S8 and Table S11).

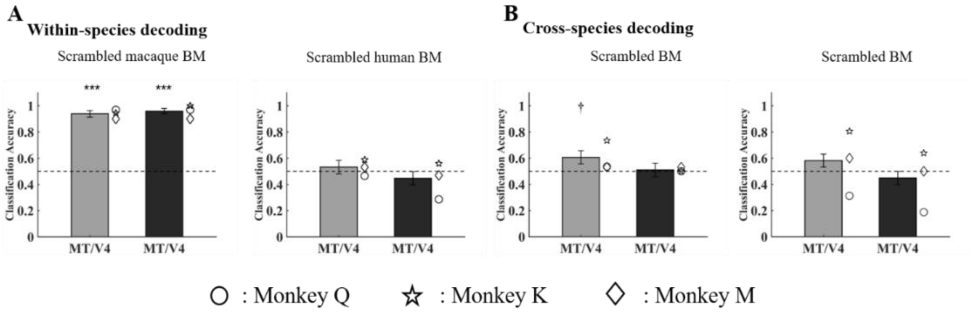

**Figure S8.** The monkey MVPA results. The classification accuracies of within-species (A) and cross-species (B) decoding of upright and inverted BM stimuli in the MT/V4. Error bars indicate standard errors. The symbols (circle, star, and diamond) on the right side of the individual bar represent results from monkeys Q, K, and M, respectively. The black dashed lines indicate chance level (50%). †, marginally significant, ***$p < 0.001$.

#### Section C: Results of combined pSTS in humans

To rule out the possibility that the species-specific representation in the human pSTS is due to that the human BM-sensitive and the macaque BM-sensitive ROIs were separately defined by two different localizers, we performed supplementary analyses on the human data.

We defined the combined pSTS using the contrast of human + macaque upright intact BM > human + macaque inverted scrambled BM (MNI coordinates: combined right pSTS, [x y z] = 50, -38,11; combined left pSTS [x y z]=-54,-44,11). First, we conducted the same ROI analysis as those in the main text on the combined pSTS. For the intact human BM, the repeated measures ANOVA revealed a significant main effect of Orientation ($F_{1,17} = 9.72$, $p = 0.006$) and a significant interaction ($F_{1,17} = 9.74$, $p = 0.006$). Post-hoc $t$-tests showed that the upright intact human BM, relative to their inverted counterparts, significantly activated the right pSTS only ($t_{17} = 3.77$, $p = 0.002$; the left pSTS, $t_{17} = 1.30$, $p = 0.211$). For the intact macaque BM, scrambled human BM, and scrambled macaque BM, no significant effects were found ($F$s < 2.04, $ps > 0.172$).

Second, we conducted the MVPA within species. For the intact human BM, significant above-chance classification of its upright and inverted version was observed in the right combined pSTS (57.8% ± 11.1; $t_{17} = 2.99$, $p = 0.033$, Bonferroni-corrected, similarly hereinafter). Similar results were found for the intact macaque BM, with classification accuracy significantly above chance level in the right combined pSTS (56.6% ± 7.8, $t_{17} = 3.59$, $p = 0.009$). For the scrambled human BM, the classification accuracy was marginally significantly higher than chance in the right combined pSTS (55.6% ± 9.2, $t_{17} = 2.56$, $p = 0.081$), while for the scrambled macaque BM, the classification accuracies were not significantly higher than chance level. Furthermore, we also conducted the MVPA across species. The results revealed that all stimuli (intact human/macaque BM and scrambled human/macaque BM) could not be well discriminated in the combined pSTS ($ts < 2.57$, $ps > 0.079$).

In addition, we conducted the DCM analysis. Once again, we found that the 'bidirectional' model had the highest posterior probability for both the intact human BM and intact macaque BM stimuli. The results showed that the modulatory effect of the upright BM stimuli was significantly stronger than that of the inverted ones on the forward connection from hMT+ to pSTS ($t_{17}$ = 2.18, $p$ = 0.043), but only for the intact human BM perception, not the intact macaque BM perception ($t_{17}$ = 1.41, $p$ = 0.176).

In summary, these supplementary analyses reaffirmed that the findings of the pSTS in the main text remained consistent, regardless of the contrast used to define the regions of interest (ROIs). The pSTS showed robust and selective activation to the human BM stimuli. In addition, the combined pSTS could decode BM within species but not across species. Finally, the functional connection between hMT+ and combined pSTS existed only for processing human BM.

#### Section D: Results of TEO in monkeys

*Definition of the TEO ROI*

To investigate whether specific subregions in the monkey STS demonstrate a species-specific processing function analogous to the human pSTS, even when they exhibit similar responses to upright and inverted BM stimuli, we utilized searchlight analyses.

First, we conducted two sets of whole-brain searchlight analyses on within-species and cross-species classifications, employing 50 voxels for each sphere (Figure S9 and S10). The classification methods were consistent with those applied in the ROI-based decoding analyses in the main text.

Subsequently, we conducted a conjunction analysis based on the two sets of searchlight results mentioned above. In accordance with the MVPA results in the human pSTS, we generated two types of maps: 1) for the within-species classification searchlight results of each type of BM stimuli, excluding scrambled humans (for which we did not detect a successful decoding cluster which was consistent across three monkeys and overlap with other types of the BM stimuli), we selected voxels with accuracies exceeding 58%; 2) for the across-species classification searchlight results (training on intact human BM stimuli and testing on intact macaque BM stimuli, and vice versa), we selected voxels with accuracies lower than 58%. The threshold of 58% was chosen based on our MVPA analysis of the MT/V4, in which we observed that an accuracy of 58% corresponded to a *p*-value of approximately 0.050 for all analyses. As shown in Figure S11, we identified a consistent cluster in the lower bank of the STS within TEO across the three monkeys.

*Univariate and MVPA Analyses on the TEO*

Next, we conducted univariate analyses on the defined TEO ROI. Unsurprisingly, our findings revealed no significant difference in response to the upright and inverted versions of all four types of BM stimuli (Figure S12, $Fs < 0.68$, $ps > 0.411$). As anticipated, TEO exhibited the ability to decode the orientation of BM stimuli, except for the scrambled human stimuli (Figure S13A, $ps < 0.001$), but was unable to perform cross-species decoding (Figure S13B, $ps > 0.471$).

*DCM Analyses in Monkeys*

To explore the functional connectivity between MT/V4 and TEO, we performed a DCM analysis akin to that conducted in humans. The DCM analysis showed that the “bidirectional modulation” had the highest posterior probability for both intact macaque and human stimuli (Figure S14A). Moreover, our results revealed significant bidirectional modulatory connections between MT/V4 and TEO for intact human and macaque BM stimuli (Figure S14B). The modulatory connections between MT/V4 and TEO for the upright stimuli were significantly higher than those for the inverted stimuli, for both intact macaque and human BM stimuli and in both forward (intact macaque: $F_{1,51} = 21.88$, $p < 0.001$; intact human: $F_{1,56} = 8.23$, $p = 0.006$) and backward (intact macaque: $F_{1,51} = 4.07$, $p = 0.049$; intact human: $F_{1,56} = 5.86$, $p = 0.019$) directions. Notably, the driving inputs of the upright stimuli were higher than those of the inverted stimuli but only for the intact macaque BM stimuli ($F_{1,51} = 8.25$, $p = 0.006$).

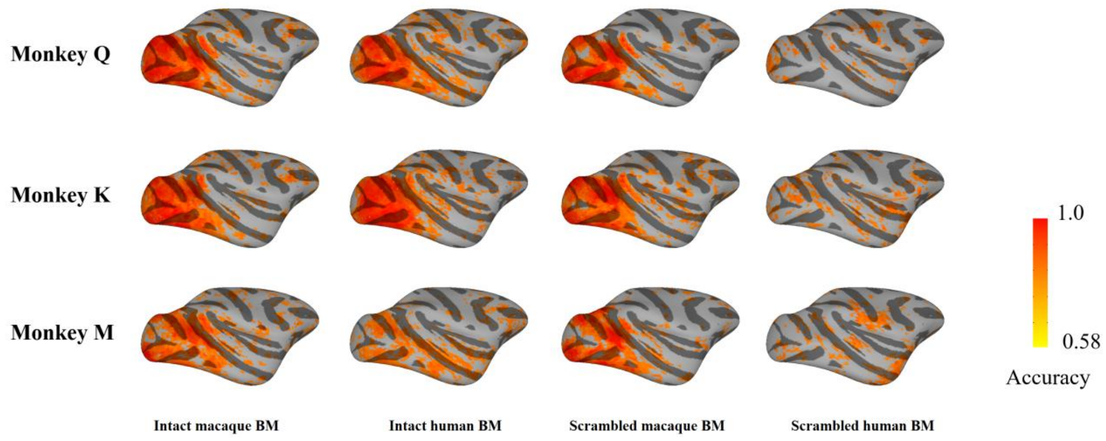

**Figure S9.** Searchlight results for the within-species classification. The color bar indicates the accuracy of BM searchlight maps.

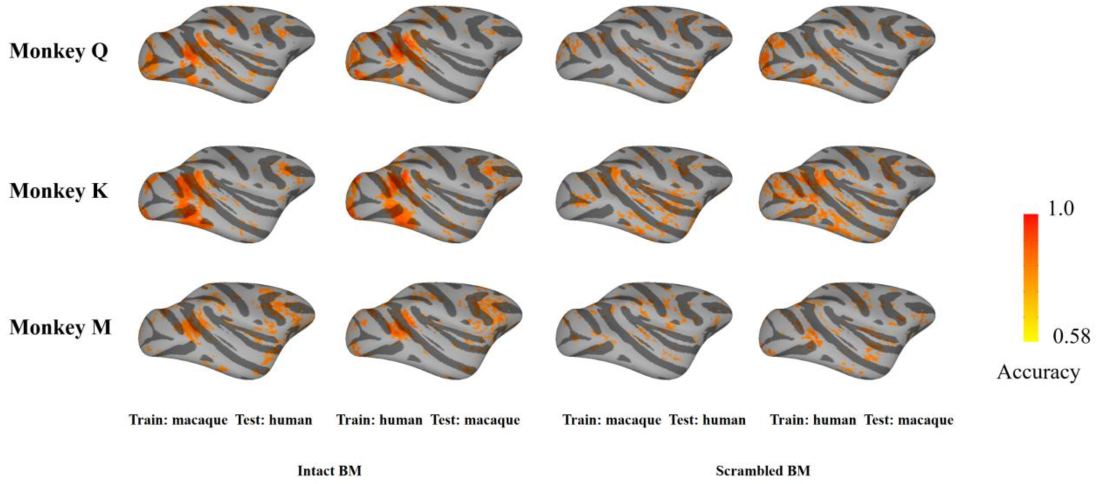

**Figure S10.** Searchlight results for the cross-species classification. The color bar indicates the accuracy of BM searchlight maps.

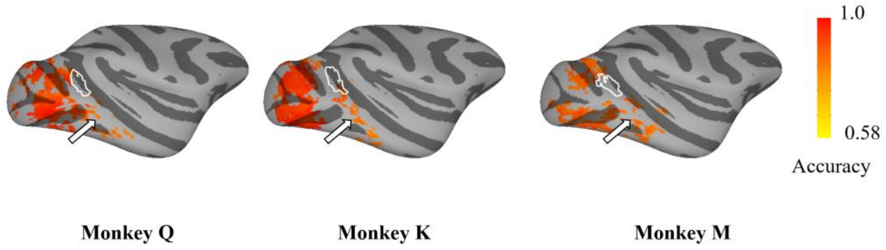

**Figure S11.** Survival voxels under the TEO definition (see section C: Definition of the TEO ROI). The color bar indicated the mean accuracy for the sum of intact macaque/human and scrambled macaque BM searchlight maps. The borders of the MT/V4 ROI are encircled by white lines. Arrows mark the approximate location of the TEO ROI.

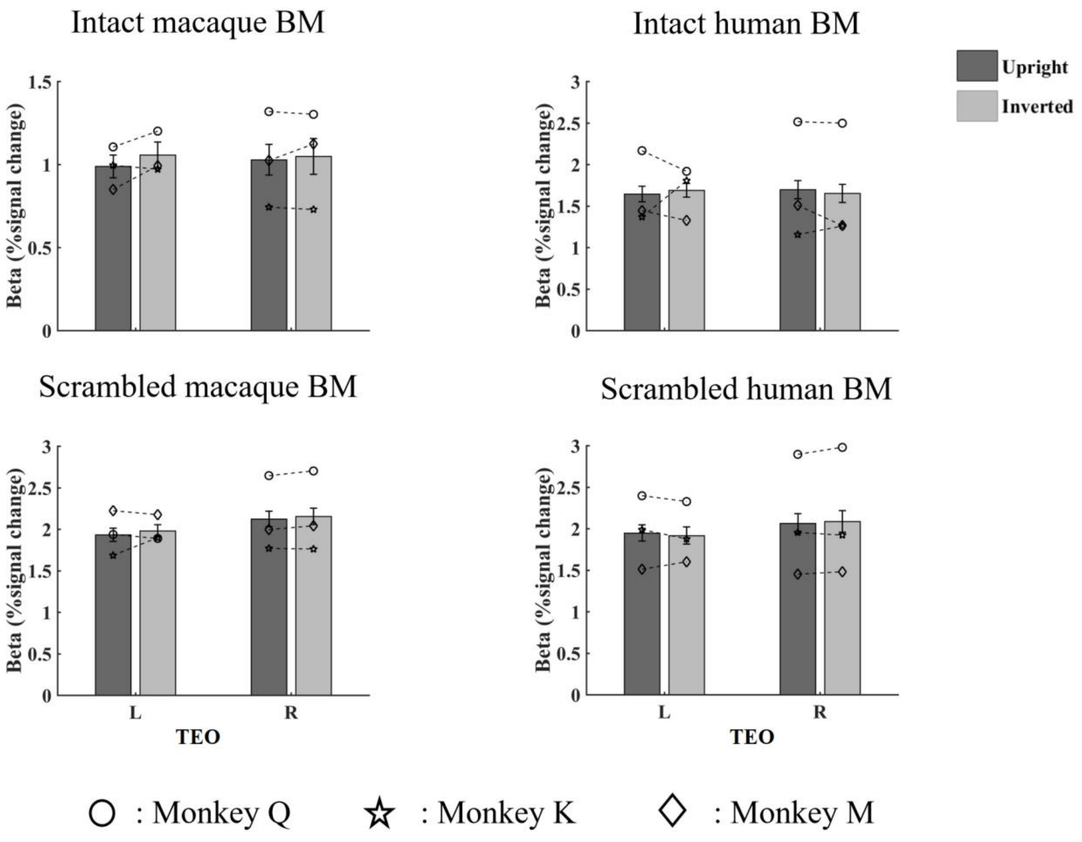

**Figure S12.** Results of the ROI analysis in the TEO. Error bars indicate standard errors of the mean. The symbols (circle, star, and diamond) within individual bars represent results from monkeys Q, K, and M, respectively.

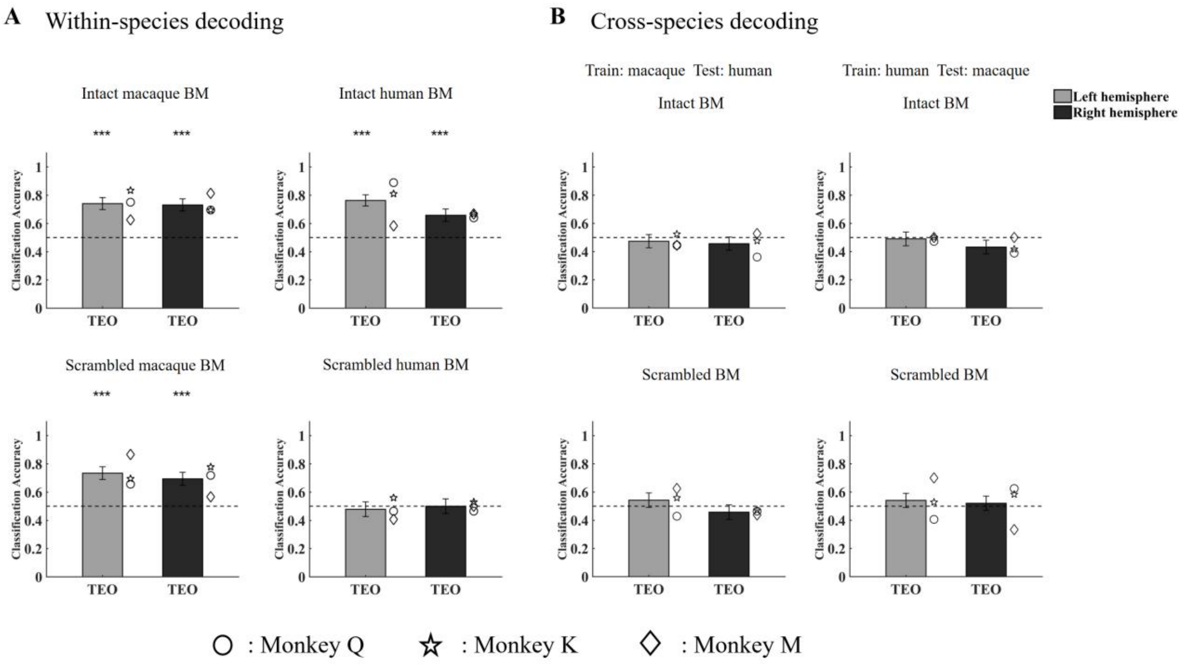

**Figure S13.** The monkey MVPA results. The classification accuracy of within-species (A) and cross-species (B) decoding of upright and inverted BM stimuli in the TEO. Error bars indicate standard errors. The symbols (circle, star, and diamond) on the right side of the individual bar represent results from monkeys Q, K, and M, respectively. The black dashed lines indicate chance level. ****p* < 0.001.

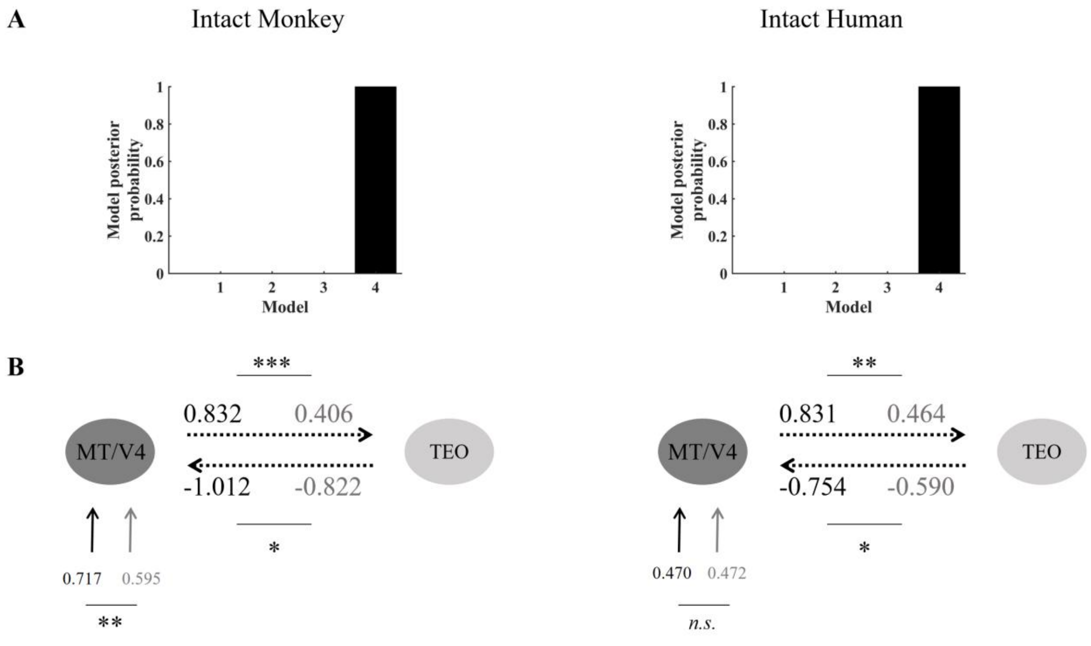

**Figure S14.** The DCM results for intact human and macaque BM perception (upright versus inverted). (A) the posterior probability for all models. (B) the parameters for driving inputs and modulatory effects (expressed in Hz). The black and the gray separately indicate the parameters from the upright and inverted BM. *n.s.,* not significant, *$p < 0.05$, **$p < 0.01$, ***$p < 0.001$.
